# Recurrent urinary tract infection and estrogen shape the taxonomic ecology and functional potential of the postmenopausal urobiome

**DOI:** 10.1101/2021.11.06.467345

**Authors:** Michael L. Neugent, Ashwani Kumar, Neha V. Hulyalkar, Kevin C. Lutz, Vivian H. Nguyen, Jorge L. Fuentes, Cong Zhang, Amber Nguyen, Belle M. Sharon, Amy Kuprasertkul, Amanda P. Arute, Tahmineh Ebrahimzadeh, Nitya Natesan, Qiwei Li, Chao Xing, Vladimir Shulaev, Philippe E. Zimmern, Kelli L. Palmer, Nicole J. De Nisco

## Abstract

Community-acquired urinary tract infection (UTI) is among the most common bacterial infections observed in humans. Postmenopausal women are a rapidly growing and underserved demographic group who are severely affected by recurrent UTI (rUTI) with a >50% recurrence rate. In this population, rUTI can persist for years, reducing quality of life and imposing a significant healthcare burden. rUTI is most often treated by antibiotics, but development of antibiotic resistance and allergy limit therapeutic options. The female urinary microbiome (urobiome) has been identified as a key component of the urogenital environment. However, compositional and functional changes in the urobiome underlying rUTI susceptibility in postmenopausal women are not well understood. Here, we used a controlled, cross-sectional cohort of postmenopausal women, to interrogate changes in urobiome structure and function linked to rUTI susceptibility by whole genome metagenomic sequencing (WGMS), advanced urine culture, estrogen metabolite profiling, and antibiotic sensitivity testing. Overall, we detected 276 bacterial, archaeal, and fungal species representing 106 genera. We find a putative commensal population consisting of species known to protect against bacterial vaginosis, such as *Lactobacillus crispatus*, within the urobiome of postmenopausal women who do not experience UTI. Integration of clinical metadata detected an almost exclusive enrichment of lactobacilli, including *L. crispatus* and *L. vaginalis*, in women taking estrogen hormone therapy (EHT). Integrating quantitative metabolite profiling of urinary estrogens with WGMS, we observed robust correlations between urobiome taxa, such as *Bifidobacterium breve* and *L. crispatus*, and urinary estrogen conjugate concentrations in women with no history of UTI that were absent in women with rUTI history. We further used functional metagenomic profiling and patient-derived isolate phenotyping to identify microbial metabolic pathways, antimicrobial resistance genes (ARGs), and clinically relevant antimicrobial resistance phenotypes enriched between disease-states. Our data indicate that distinct urobiome metabolic and ARG signatures are associated with current rUTI status and history. Importantly, we observed that rUTI history leaves an imprint of enriched ARGs even in women not currently experiencing UTI. Taken together, our data suggests that rUTI history and estrogen strongly shape the functional and taxonomic composition of the urobiome in postmenopausal women.

## Introduction

Urinary tract infection (UTI) is among the most common adult bacterial infections and imparts a particularly significant medical burden on women, with more than 50% of women suffering UTI in their lifetime (Gaitonde et al., 2019; Jhang and Kuo, 2017). Historically, UTI has largely been underprioritized in medical research due to low mortality rates and the effectiveness of available antibiotics. However, UTI is a disease of disproportionate burden as age is one of the strongest associated risk factors for UTI and the development of recurrent UTI (rUTI) (Flores-Mireles et al., 2015). Indeed, approximately 50% of UTIs in postmenopausal (PM) women develop into rUTI, which is clinically defined as ≥3 symptomatic UTIs in 12 months (Gaitonde et al., 2019; Malik et al., 2018b). rUTI can last for years, dramatically decreasing quality of life and, if treatment is unsuccessful, can develop into life-threatening urosepsis. Current therapeutic strategies rely on the use of antibiotics to achieve urinary tract sterility (Flores-Mireles et al., 2015; Malik et al., 2018a; Neugent et al., 2020). However, increasing rates of antibiotic refractory rUTI make this strategy unsustainable and ultimately ineffective (Malik et al., 2018a). Alternate therapeutic strategies are therefore needed to increase quality of life and reduce adverse outcomes for women with rUTI.

A promising source of therapeutic strategies for rUTI lies in modulating or restoring the urinary microbiome, termed here the “urobiome” (Stamm and Norrby, 2001; Wolfe and Brubaker, 2019). Decades of medical dogma have largely assumed sterility of urine and the urinary tract; however, a large body of work has robustly established the existence of a human urobiome (Brubaker and Wolfe, 2017; Hilt et al., 2014; Lewis et al., 2013; Price et al., 2019; Siddiqui et al., 2011; Wolfe et al., 2012). Initial taxonomic analyses have associated urobiome compositional dysbiosis with urinary incontinence, overactive bladder, and bladder cancer (Bucevic Popovic et al., 2018; Karstens et al., 2016; Pearce et al., 2014). Still, fundamental knowledge of urobiome composition and function in PM women, especially in the context of rUTI susceptibility, is lacking. Studies surveying the microbial ecologies of the urobiome associated with UTI have almost exclusively focused on premenopausal women or cohorts of mixed age (Barraud et al., 2019; Brubaker and Wolfe, 2017; Hilt et al., 2014; Neugent et al., 2020; Price et al., 2019; Siddiqui et al., 2011; Wolfe et al., 2012). As a result, the relationship between the urobiome and rUTI susceptibility is poorly understood in PM women. A 2021 report showed that premenopausal and PM women displayed different core urinary microbiota at the genus level, providing strong rationale to characterize the PM urobiome in urogenital disease (Ammitzboll et al., 2021).

The female urobiome is reported to be interconnected with the vaginal microbiome (Thomas-White et al., 2018). For example, D(-)Lactate-producing lactobacilli, known to protect the vagina from colonization by bacterial and fungal pathogens, have been consistently observed in the female urobiome in multiple independent studies (Edwards et al., 2019; Pearce et al., 2014). These observations beg the question of whether these known protective vaginal species serve a similar role in the urobiome. A 2011 clinical trial found a moderate reduction of rUTI incidence among women receiving an intravaginal *L. crispatus* probiotic (Stapleton et al., 2011). While this study was performed in premenopausal women and has yet to be confirmed in a larger clinical study, it does suggest that lactobacilli may support urinary tract health.

To date, no whole genome metagenomic sequencing (WGMS) datasets have been generated to profile the urobiome of PM women. Given the genomic diversity observed within and among taxonomic clades, metagenomic information beyond 16S rRNA sequence enrichment is needed to assess the functional potential of microbial communities (Quince et al., 2017). Whole-metagenome analysis of the urobiome is required to identify the genes and metabolic pathways associated with urinary tract health. Here, we present a WGMS survey of the urobiome of a strictly curated, cross-sectional cohort of PM women separated into three groups defined by rUTI history and current UTI status. Taxonomic biomarker analysis detected a microbial signature that suggests an imprint of past UTI remains in the urobiome. We also observed a striking association between the use of estrogen hormone therapy (EHT) and the presence of vaginal lactobacilli in the urobiome. Further, we observed strong *Lactobacillus* enrichment among women using oral and patch EHT, but not women using vaginal EHT. These findings mirror the detected concentrations of excreted urinary estrogen metabolites in women using EHT, which we found positively correlated with abundances of potentially protective urinary taxa in women with no UTI history. Finally, we found that the resistome of the urinary microbiota is dramatically altered in women with rUTI history even in the absence of active infection. Taken together, these results suggest that both urobiome taxonomy and functional potential are shaped by rUTI history and EHT in PM women.

## Results

### Cohort curation, metagenomic DNA preparation, and whole genome metagenomic dataset generation

rUTI often follows a cyclic pattern of infection (relapse) interrupted by periods of remission (Figure 1A). To model this pattern of relapse and remission, PM women were striated into three groups based on rUTI history. Group 1 served as a healthy comparator and consisted of PM women with no lifetime history of symptomatic UTI (No UTI History), group 2 consisted of PM women with a recent history of rUTI but no active UTI at the time of urine donation (rUTI Remission), and group 3 consisted of PM women with a history of rUTI and an active, symptomatic UTI at the time of urine donation (rUTI Relapse) (Figure 1B). All recruited women passed strict inclusion criteria for uncomplicated rUTI. We determined that 25 women per group were sufficient to balance *a-priori* sample size estimation (Figure S 1A, B) with clinical feasibility and enrollment rates. The final cohort was balanced for sample size, Race, BMI, smoking history, EHT use, urine pH, and urinary creatinine concentration. It should be noted the women in the rUTI Relapse group tended to be older with a median age of 76 compared to 67 in the No UTI History and 68 in the rUTI Remission groups (Table S1).

**Figure 1.**
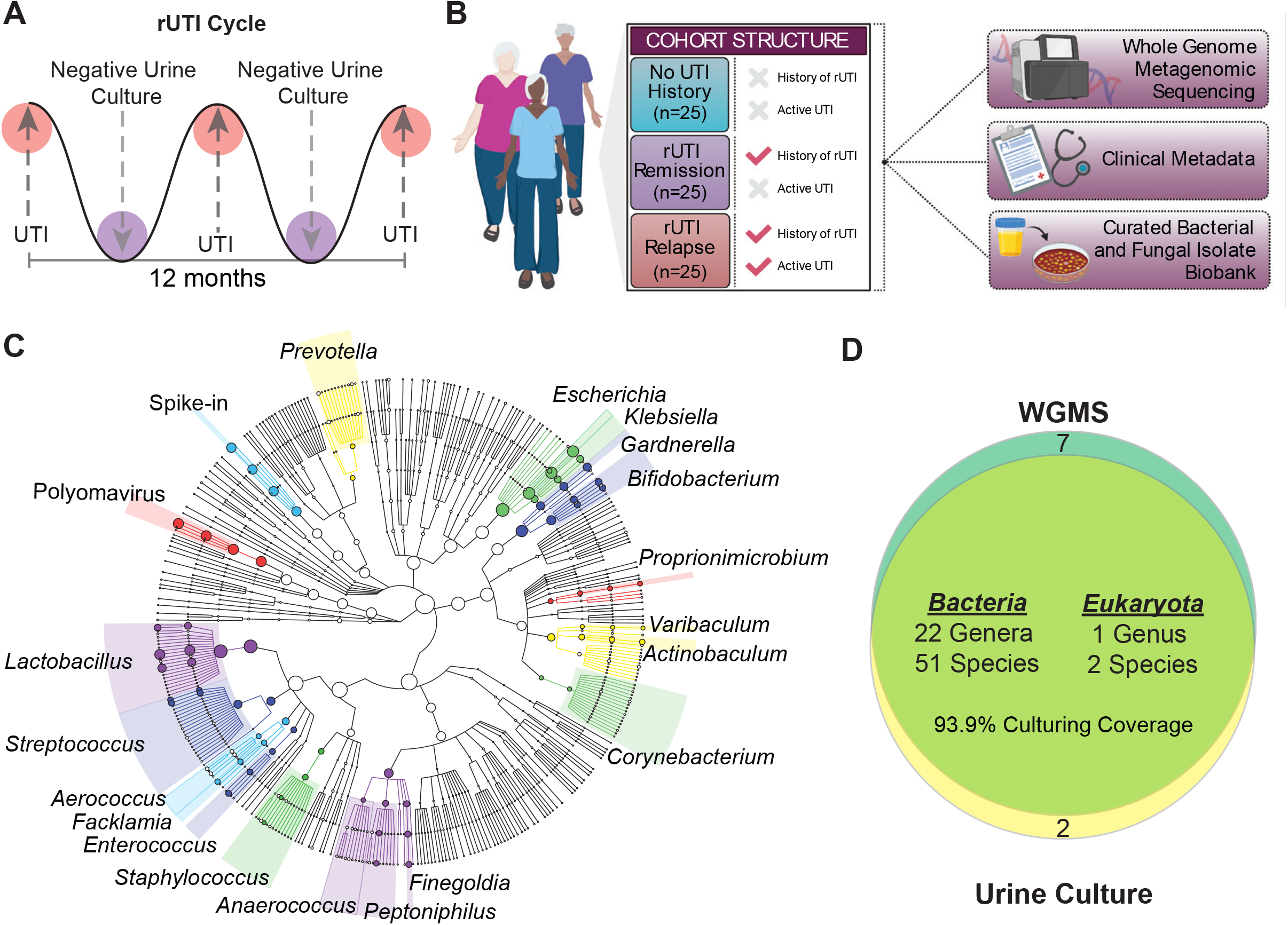
Study design and summary of genera detected by WGMS and advanced urine culture. (A) Illustration of rUTI cycle depicting periods of active, symptomatic UTI with positive urine culture followed by periods of remission with negative urine culture. (B) Diagram of cohort structure and datasets generated for the study created with BioRender.com. (C) Taxonomic cladogram of top 20 genera detected in all metagenomes (*n*=75) by Metaphlan2. Node size indicates relative abundance and branch length is arbitrary. (D) Venn diagram depicting the coverage of advanced urine culture calculated at the genus level considering all bacterial genera with >5% WGMS relative abundance in at least one patient.

Urine was collected via the clean-catch midstream method and is therefore representative of the urogenital microbiome (Karstens et al., 2018), which is includes the bladder, urethral, and, in some cases, vaginal microbiomes. Metagenomic DNA yields reflected the anticipated biomass of the urobiome in each group with the highest DNA yields observed in the rUTI Relapse group (Figure S 1C). We observed an average of 67.6% host contamination within the WGMS data (Figure S 1D). Previous reports of human contamination in WGMS sequencing of the urobiome range from 1-99% of obtained reads (Moustafa et al., 2018; Quince et al., 2017). To measure potential background and environmental taxonomic signals, a pure water sample was randomly inserted into the metagenomic DNA isolation and sequencing workflow (Segata et al., 2012). Most microbial reads observed in the water control mapped to common nucleic acid purification kit and environmental contaminants (Figure S 2A) (Salter et al., 2014). Except for known members of the human microbiome, these background taxa were censored from the data.

### Validation of viable urobiome species through advanced urine culture and WGMS hybrid taxonomic profiling

To validate the presence of living microbiota within the urobiomes sampled, we coupled WGMS with advanced urine culture, a modification of the previously reported enhanced quantitative urine culture protocol (Price et al., 2016). Taxonomic profiling by WGMS detected 276 bacterial, archaeal, and fungal species across 106 genera within the aggregate urobiome of all three groups. The sampled urobiomes were dominated by the kingdom, Bacteria, which represented 99.4% of the detected non-viral, microbial taxa (Figure 1C). Consistent with the observed taxonomic composition of urobiomes studied to date (Brubaker and Wolfe, 2017; Siddiqui et al., 2011), the detected bacterial taxa across the cohort belonged to four major phyla: Firmicutes (44.7%), Actinobacteria (22.3%), Proteobacteria (20.6%), and Bacteroidetes (12%) (Figure 1C). Advanced urine culture captured 93.9% of bacterial genera detected in WGMS with observed aggregate relative abundance ≥5% in any sample (Figure 1D). Patient-level culture coverage is reported in Figure S 2A. The most frequent cultivable genera across all samples were *Lactobacillus, Escherichia, Streptococcus, Bifidobacterium, Gardnerella, Klebsiella, Staphylococcus, Finegoldia, Enterococcus*, and *Facklamia*. Pure isolates from every cultivable species were assembled into a biobank of 896 bacterial and fungal isolates.

### rUTI History is not associated with large-scale alterations of urobiome ecological structure in the absence of active infection

We next analyzed the genus and species-level taxonomic profiles within the No UTI History, rUTI Remission and rUTI Relapse groups (Figure 2A, B, Figure S 3A). rUTI Relapse group urobiomes were mainly dominated by single bacterial uropathogens with little detected abundance of Fungi and *Archaea* (Figure 2B, Figure S 3A). The most prevalent bacterium was *Escherichia coli* (15/25, 60%), the major uropathogen among most types of UTI (Flores-Mireles et al., 2015). Along with *E. coli*, we detected known uropathogens, *Klebsiella pneumoniae* (2/25, 8%), *Enterococcus faecalis* (1/25, 4%), and *Streptococcus agalactiae* (1/25, 4%). We observed a low relative abundance of fungal species within the rUTI Relapse urobiomes including *Candida glabrata* and *Malassenzia globosa*. Similarly low relative abundances of archaeal taxa were detected, such as, *Methanobrevibacter spp*. (Figure S 3A). The most observed viral taxa were JC polyomavirus (4/25 16%) but Human herpes virus 4 (1/25 4%) and Enterobacteria phage lke (1/25 4%) were also detected (Figure S 3B).

**Figure 2.**
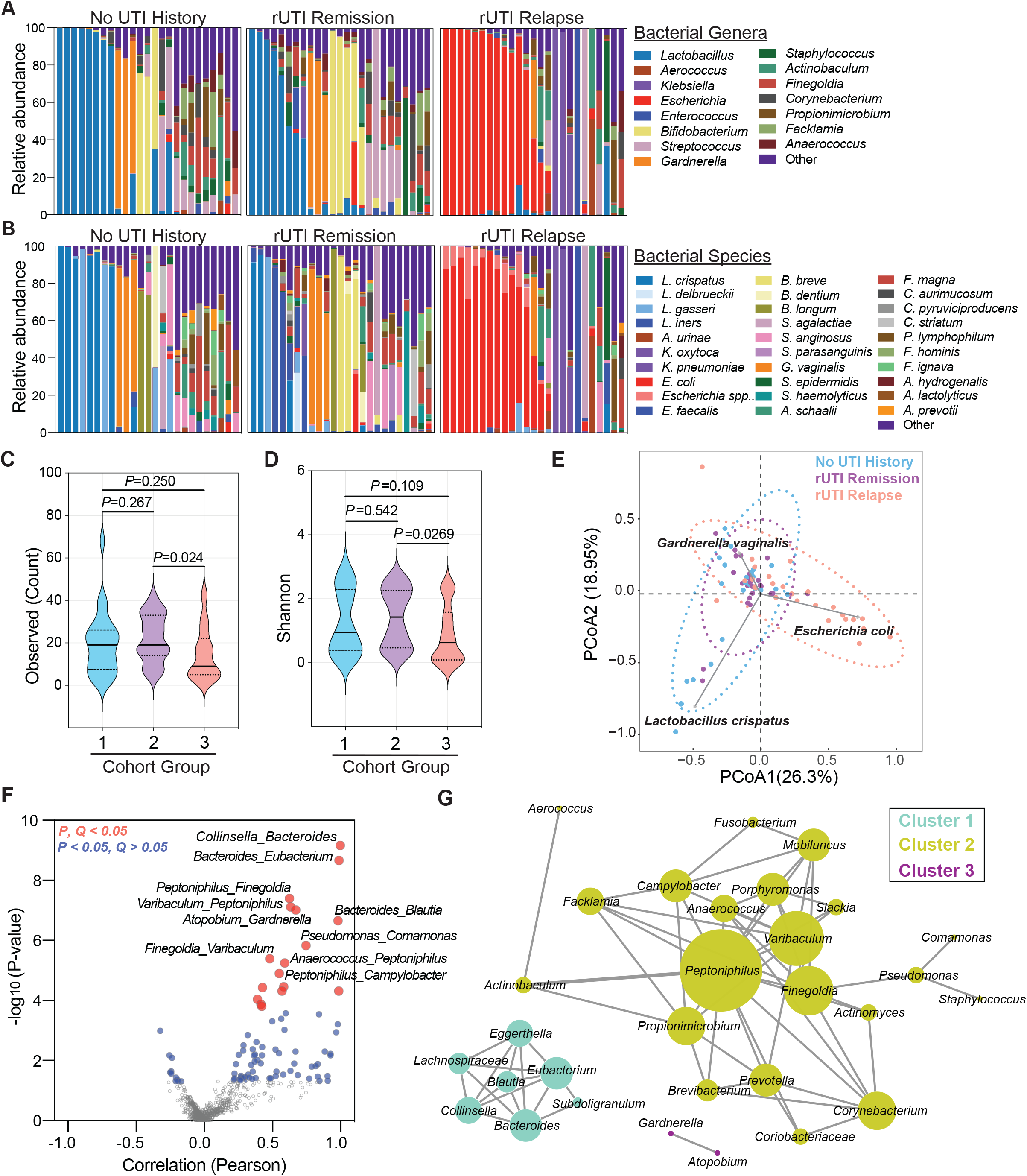
The bacterial taxonomic profile of rUTI in PM women. Genus (A) and species-level (B) taxonomic profiles of the top 15 bacterial genera among cohort groups (No UTI History (*n*=25), rUTI Remission (*n*=25), rUTI Relapse (*n*=25)). Remaining genera or species are combined into “Other”. Alpha-diversity comparison of observed species counts (C) and Shannon index (D) between groups (1 = No UTI History, 2 = rUTI Remission, 3 = rUTI Relapse). Solid lines represent medians while dotted lines represent the interquartile range. *P*-value generated by Kruskal-Wallis test with Dunn’s multiple correction post hoc. (E) Beta-diversity of the WGMS data by DPCoA. Each dot represents an individual sample color-coded by group (No UTI history in blue, rUTI remission in purple, and rUTI relapse in salmon). Vectors (grey) represent major discriminating loadings (*i*.*e*. species). (F) Volcano plot depicting co-occurrence of genera within the WGMS dataset by Pearson correlation. *P*-value generated by permutation. Red dots represent associations with an FDR-corrected *P*-value < 0.05. Blue dots represent associations with a nominal *P*-value < 0.05, but an FDR-corrected *P*-value > 0.05. (G) Network analysis of all genus-level co-occurrences with nominal *P*-value < 0.05. Nodes represent genera. Edges are defined by Pearson correlation and node size is proportional to the degree, or connectivity, of the node.

The most frequently observed bacterial species in urobiomes of women without active UTI (No UTI History and rUTI Remission) belonged to the genera *Lactobacillus, Bifidobacterium, Gardnerella, Streptococcus, Staphylococcus*, and *Actinobaculum* (Figure 2 A, B). A subset of these samples (54%) was dominated by one taxon while others were diverse and exhibited no single dominant taxon. We observed 13 patients (24%) in the No UTI History and rUTI Remission groups with a >50% relative abundance of various *Lactobacillus spp*., including *L. crispatus, L. iners*, and *L. gasseri* (Figure 2B). Interestingly, we also observed a subset of urobiomes in the No UTI History and rUTI Remission groups dominated by *Bifidobacterium spp*., such as *B. breve, B. dentium*, and *B. longum*, as well as by *Gardnerella vaginalis* (Figure 2B). Fungal and archaeal species were observed in low abundance (0-8.8% relative abundance) in the No UTI History and rUTI Remission urobiomes and included *Candida albicans, C. glabrata*, and *Candida dubliniensis*, as well as *M. globosa, Naumovozyma spp*., and *Eremothecium spp* (Figure S 3A). Observed archaeal species within the No UTI History and rUTI Remission urobiomes included *Methanosphaera stadtmanae* and *Methanobrevibacter spp*. Viral taxa were more frequently observed in the No UTI history and rUTI Remission groups than in the rUTI Relapse group and included JC, BK, and Merkel cell polyomaviruses (Figure S 3B).

To model the urobiome ecological structure within the three groups, we calculated alpha-diversity indices including the observed taxa count, Shannon, Simpson, Chao 1, and ACE indices. Women in the No UTI History and rUTI Remission groups had similarly diverse urobiomes across different indices (Figure 2C, D, Figure S 4 A, B, C). These data suggest that if there are differences in the urobiomes of PM women who are susceptible to rUTI (rUTI Remission) versus those who are not (No UTI History), they are not reflected in alpha-diversity metrics. The rUTI Relapse group exhibited significantly lower alpha-diversity as compared to the rUTI Remission cohort (Figure 2C, D, Figure S 4 A, B, C).

To assess large-scale taxonomic signatures associated with rUTI, we used double principal coordinate analysis (DPCoA) (Pavoine et al., 2004). Visualization of the first two PCoAs revealed that the urobiomes of the rUTI Relapse group clustered along a vector defined by *E. coli* and were ecologically distinct from the urobiomes of the No UTI History and rUTI Remission groups. The No UTI History and rUTI Remission groups exhibited relatively similar clustering patterns in the first two PCoAs (Figure 2E), and clustered along opposing vectors defined by the enrichment of either *L. crispatus* or *G. vaginalis*, which are associated with a healthy vaginal microbiome or bacterial vaginosis, respectively (Ravel and Brotman, 2016; Ravel et al., 2011). This similar clustering of the No UTI history and rUTI Remission cohorts suggests that a history of rUTI does not significantly alter the large-scale taxonomic structure of the urobiome in PM women.

### Taxonomic profile of the PM urobiome displays co-occurrence structure

Microbial communities can harbor intricate interactions between member taxa (Alteri et al., 2015; Keogh et al., 2016). Exceedingly little is known about interactions and co-occurrence of bacterial species within the urobiome. We performed taxonomic association analysis to determine the co-occurrence structure of the PM urobiome and identified 87 statistically significant genus-level associations (*P*<0.05) (Figure 2F). After multiple hypothesis testing correction, a total of 17 unique associations exhibited robust statistical significance (*Q*<0.05) (Figure 2F). Network visualization of significant positive associations (*P*<0.05) revealed 3 non-interacting microbial clusters (Figure 2G). Cluster 1 member taxa included genera previously associated with vaginal infections, such as *Bacteroides* and *Blautia* (Ceccarani et al., 2019). Cluster 2 exhibited the largest member set and diversity, and captured associations between genera known to inhabit the urobiome (i.e. *Peptoniphilus* and *Finegoldia*) but whose association has not yet been reported (Anglim et al., 2021; Thomas-White et al., 2017). Cluster 2 grouped strongly around the genus *Peptoniphilus*. Cluster 3 was identified as a pairwise interaction between the genera *Gardnerella* and *Atopobium*, two taxa that are associated in vaginal dysbiosis and bacterial vaginosis (Bradshaw et al., 2006; Hardy et al., 2016; Hardy et al., 2015). We also observed anticorrelated taxa (Figure 2 F, Figure S 4 D). The two main hubs of the negative correlation network were the genera *Lactobacillus* and *Escherichia*. Of note, the most significant negative association was observed between the *Lactobacillus* and *Peptoniphilus* (Figure 2 F, Figure S 4 D). Taken together, these data define patterns of co-occurrence within the urobiome and suggest candidate taxa, (i.e. *Lactobacillus, Peptoniphilus*, and *Escherichia)* that may act as hubs of community structure.

### Taxonomic biomarker analysis reveals that rUTI history alters the species-level taxonomic signature of the urobiome

Although we did not detect large scale taxonomic differences between the rUTI Remission and No UTI History groups, we hypothesized that small scale differences (e.g. enrichment of a single genus or species) may contribute to differential rUTI susceptibility between groups. We therefore performed genus and species-level differential taxonomic enrichment analysis between the No UTI History and rUTI Remission urobiomes using both linear discriminant analysis of effect size (LEfSe) and a Bayesian microbial differential abundance (BMDA) model (Li et al., 2019; Segata et al., 2011). While LEfSe performs a non-parametric assessment of differential abundance, BMDA can account for sparsity, over-dispersion, and uneven sampling depth. Though LEfSe detected no differentially abundant taxa between the No UTI History and rUTI Remission groups, BMDA detected multiple differentially abundant taxa. BMDA detected two genera, *Aerococcus* and *Lactobacillus*, as well as two species of lactobacilli, *L. vaginalis* and *L. crispatus*, as enriched in the No UTI History group (Figure 3A, B). At the genus-level, *Klebsiella, Gemella, Bacteroides, Clostridiales Family XIII Incertae Sedis unclassified, Eggerthella*, and *Escherichia* were among the most significantly enriched in the rUTI Remission group. Fifteen species were identified as significantly enriched in the rUTI Remission group, including *Ureaplasma parvum, Bacteroides uniformis, Brevibacterium massiliense, Anaerococcus hydrogenalis, Actinomyces turicensis, Prevotella timonensis, E. faecalis, Staphylococcus hominis, Peptoniphilus lacrimalis, Corynebacterium pseudogenitalium, Actinomyces europeaus, Facklamia hominis, Finegoldia magna, Anaerococcus prevotii*, and *Staphylococcus epidermidis* (Figure 3B). Many of these taxa, such as the genera *Bacteroides, Streptococcus, Escherichia, Ureaplasma, Finegoldia*, and *Gemella*, have been found in the vaginal microbiome during infection (Ceccarani et al., 2019; Shipitsyna et al., 2013). Furthermore, *A. turicensis* and *A. europaeus* are known to be associated with UTI (Kononen and Wade, 2015). Taken together, these results suggest that rUTI history may leave an imprint on urobiome composition that may be missed by common ecological indices (alpha or beta diversity) and differential abundance pipelines that do not consider the sparsity, over-dispersion and uneven sampling that is common in low biomass microbiomes like the urobiome (Karstens et al., 2018).

**Figure 3.**
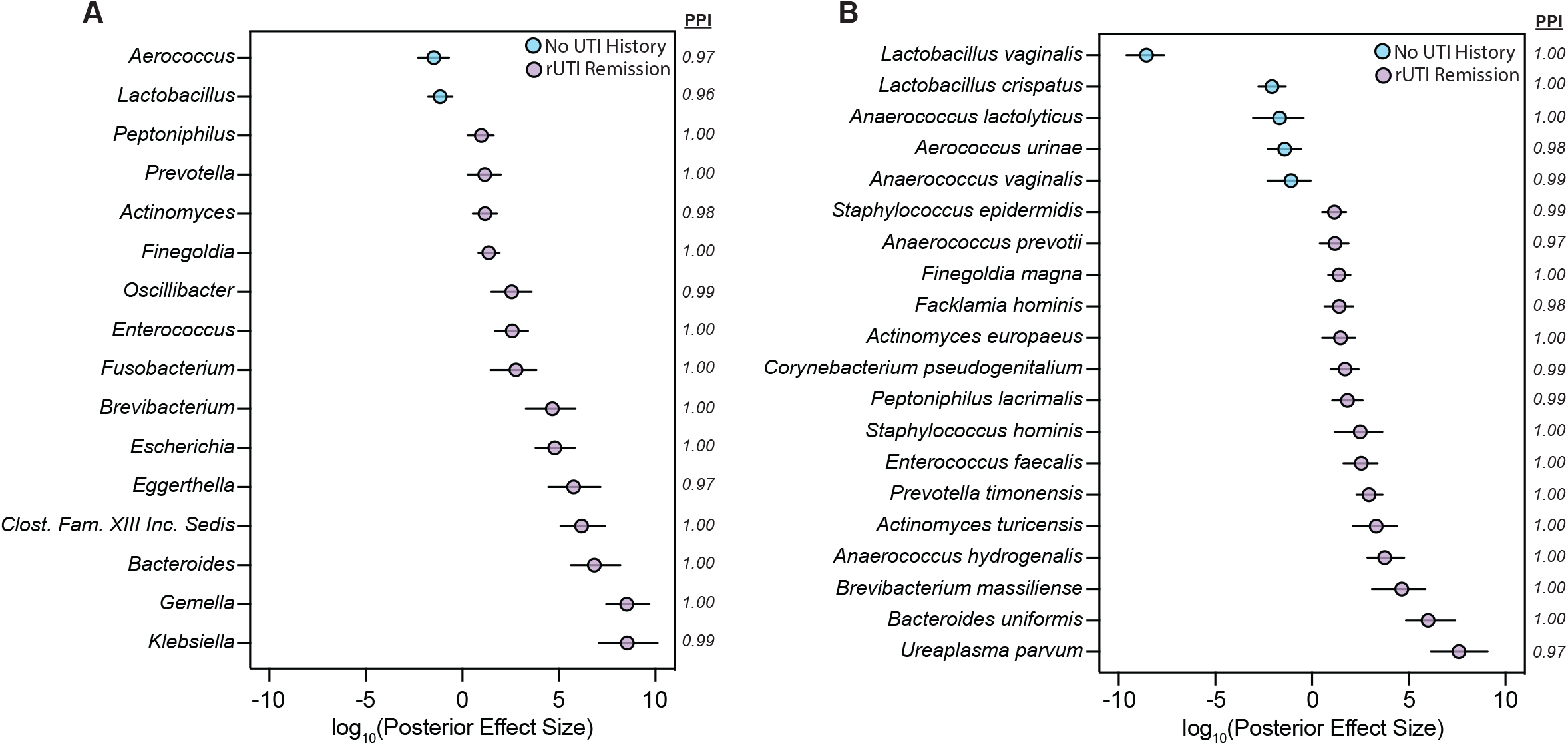
Bayesian modeling detects the taxonomic imprint of rUTI history on the urobiome of PM women. Bayesian differential abundance model comparing genus (A) and species-level (B) taxonomic enrichment between the No UTI history (*n*=25) and rUTI Remission (*n*=25) groups. Dots, indicating the log_10_(posterior effect size), are color-coded by group enrichment (No UTI history in blue and rUTI Remission in purple). Lines indicate the 95% credible interval and PPI denotes posterior probability index.

### Urobiome taxonomic structure differs in women using estrogen hormone therapy

Given that many of the urobiomes of women without active rUTI were dominated by species of lactobacilli (26%, 13/50) (Figure 2 A, B), we sought to further characterize this taxonomic enrichment in the No UTI History and rUTI Remission groups. We screened the cohort-associated clinical metadata for variables associated with *Lactobacillus* abundance. Interestingly, we found that estrogen hormone therapy (EHT) was strongly associated with the presence of *Lactobacillus* in the urobiome (Figure 4 A, B, C). Ecological modeling revealed that the urobiomes of EHT(+) women were significantly less diverse than those of EHT(-) women and tended to be dominated by a single species of lactobacilli (Figure 4 D, E, F). To identify taxa associated with EHT use, we performed differential taxonomic enrichment analysis using LEfSe and BMDA. Notably, LEfSe found an enrichment of the genus *Lactobacillus* in EHT(+) women and the genus *Streptococcus*, and EHT(-) women (Figure 4 G). BMDA captured a similar result but further resolved species-level differential enrichment (Figure 4 H). *L. crispatus* and *L. vaginalis*, were significantly enriched in the urobiomes of EHT(+) women and *Streptococcus mitis/oralis/pneumoniae* (*S. m/o/p*) group, *Streptococcus infantis*, and *Atopobium vaginae* were enriched in the EHT(-) group (Figure 4H).

**Figure 4.**
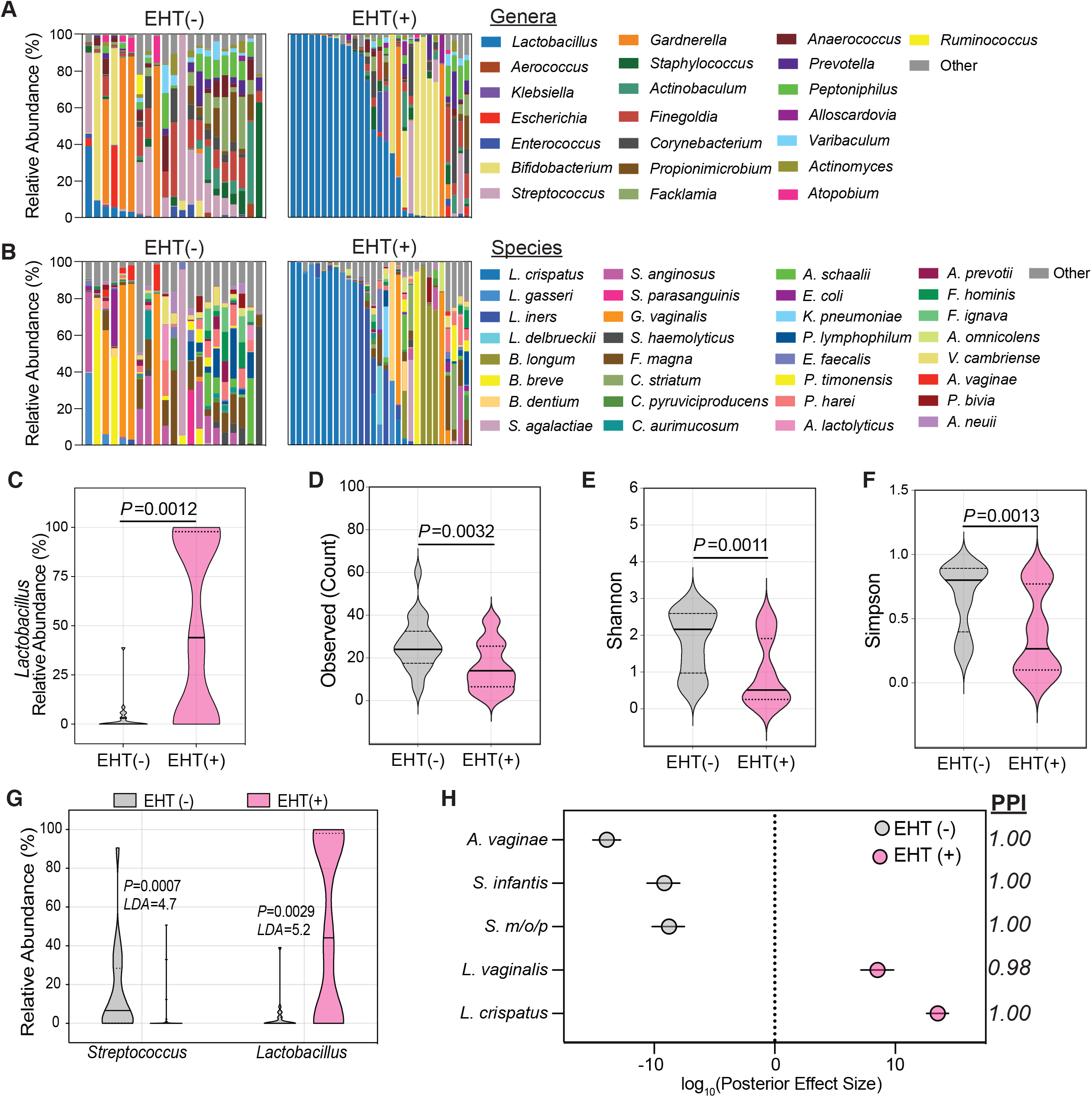
Estrogen hormone therapy shapes the urobiome of PM women. Genus (A) and species-level (B) taxonomic profiles of the relative abundance of the top 22 bacterial genera among EHT(-) (*n*=21) and EHT(+) (*n*=29) women in the No UTI History and rUTI Remission groups. All genera or species not within the top 22 are combined into “Other”. (C) Comparison of *Lactobacillus* relative abundance between EHT(-) (grey) and EHT(+) (pink) women in the No UTI history and rUTI Remission groups. Observed species count (D), Shannon index (E), and Simpson index (F) comparisons between EHT(-) (grey) and EHT(+) (pink) women in the No UTI history and rUTI remission groups. Violin plots depict the smoothed distribution of the data. Solid lines represent medians while dotted lines represent the interquartile range. *P*-values generated by Wilcoxon rank-sum. (G) Two significantly differentially enriched genera (LDA > 4.5) detected by LEfSe between EHT(-) (grey) and EHT(+) (pink) women. LDA denotes the log_10_(linear discriminant analysis score) and the *P-*value was generated by LEfSe. (H) Differentially enriched taxa between EHT(-) (grey) and EHT(+) (pink) women in the No UTI history and rUTI Remission cohorts detected by BMDA. Dots indicate log_10_(posterior effect size) and PPI denotes posterior probability index. S. m/o/p denotes *Streptococcus mitus/oralis/pneumoniae*.

### Urinary estrogen concentration is positively correlated with urobiome *Lactobacillus* abundance in PM women with No UTI history

EHT can be administered via multiple modalities including oral supplementation, transdermal patch, and topical vaginal cream (Lobo, 2017). Separating women by EHT modality, we observed that women using oral and patch EHT had significant urobiome enrichment of *Lactobacillus* while urobiome *Lactobacillus* enrichment varied widely in women using vaginal EHT (vEHT) and was not significantly different from EHT(-) women (Figure 5A). We hypothesized that vEHT may differ from oral and transdermal patch in composition, metabolism, or dosage. We therefore optimized a published targeted liquid-chromatography mass spectrometry (LC-MS) method to quantify excreted urinary estrogen conjugates of women in the No UTI History and rUTI Remission groups (van der Berg et al., 2020). Limiting our analysis to the known major excreted sulfate and glucuronide conjugates of estrone (E1) and 17β–estradiol (E2), we observed significantly higher urinary concentrations of E1 and E2 sulfates and glucuronides in women using oral EHT. This observation was concordant with urinary *Lactobacillus* abundance (Figure 5 B-D Figure S 6 A-D). Women using patch EHT also exhibited high urinary *Lactobacillus* abundance and a significant enrichment of urinary E1-sulfate compared to EHT(-) women (Figure S 5C). Consistent with urinary *Lactobacillus* abundance, we observed no statistically significant difference in urinary estrogen conjugate concentrations between vEHT(+) women and EHT(-) women (Figure 5 A-D Figure S 6 A-D).

**Figure 5.**
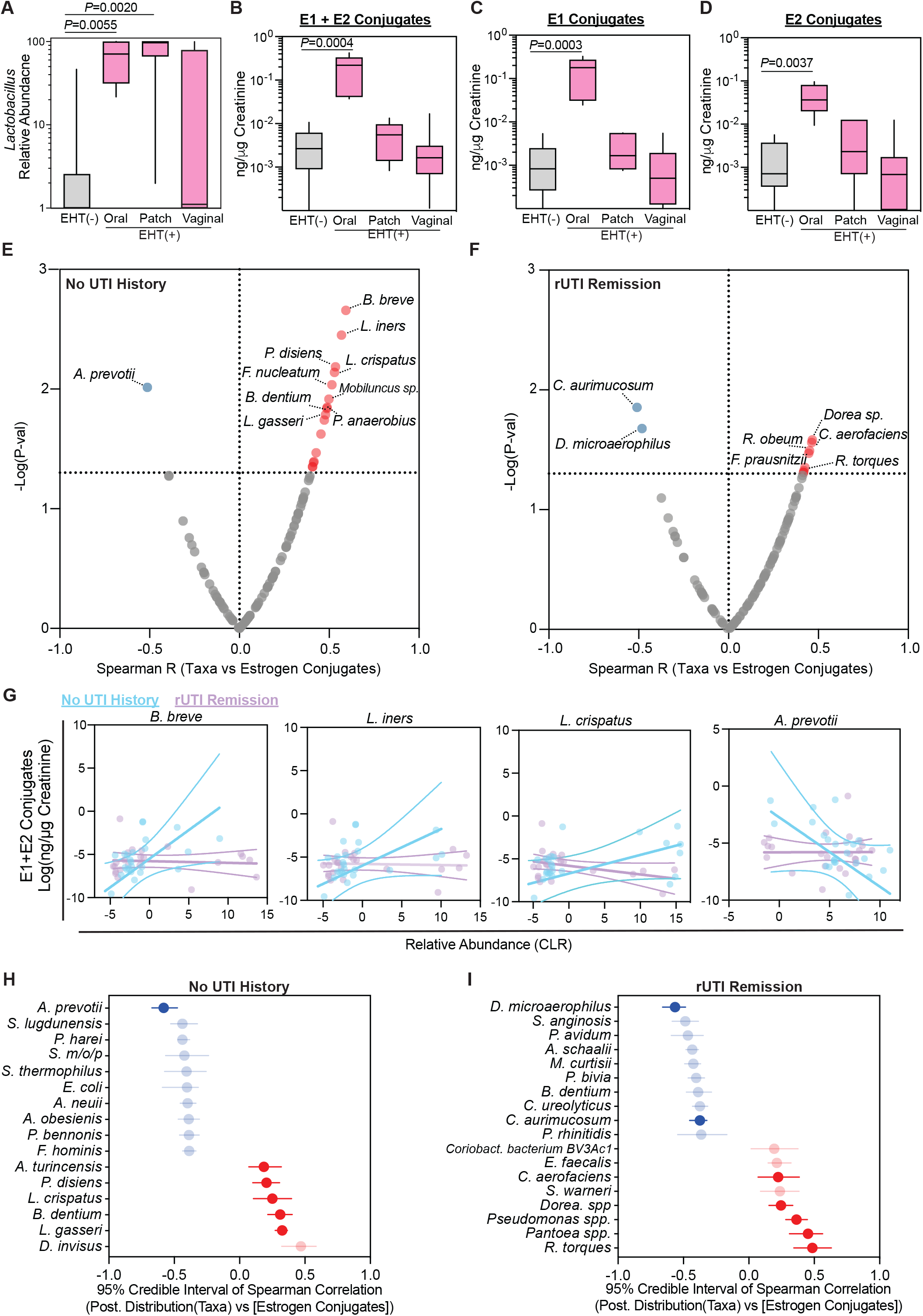
Distinct taxa-urinary estrogen metabolite associations between PM women with and without rUTI history. (A) Comparison of the relative *Lactobacillus* abundance between EHT(-) (grey, *n*=21) and EHT(+) (pink, *n*=29) women separated by EHT modality (Oral, Patch, Vaginal). Creatinine (Cr)-normalized summed estrogen conjugates (B), E1 conjugates (C) and E2 conjugates (D) measured in No UTI History and rUTI Remission urine separated by EHT modality (EHT(-) (n=20), Oral (*n*=6), Patch (*n*=6), Vaginal (*n*=17)). For all box and whisker plots, error bars are drawn from minimum to maximum, boxes represent interquartile range, solid lines denote the median, and *P*-value generated by Kruskal-Wallis test with Dunn’s multiple correction post hoc. Spearman correlation of bacterial species with summed Cr-normalized urinary estrogen conjugates in No UTI History (E) and rUTI remission (F) groups. *P*-value generated by permutation. Red and blue dots represent significant (*p<*0.05) positive and negative associations, respectively. (G) Correlation scatter plots of *B. breve, L. iners, L. crispatus*, and *A. prevotti*, and summed Cr-normalized urinary estrogen conjugate associations among No UTI History (*n*=25) (blue) and rUTI Remission women (*n*=23) (purple). Linear regression trend line (solid line) is shown with 95% confidence intervals. Bayesian correlation point estimates and 95% credible interval of posterior correlation (Spearman) for the top 10 taxa and Cr-normalized summed urinary estrogen conjugates in the (H) No UTI History and (I) rUTI Remission groups. Blue indicates negative while red indicates positive correlation. Significant correlations also found in the non-Bayesian analysis are bolded.

We further sought to determine if rUTI history differentially affected EHT-associated taxa by dichotomizing patients by No UTI History and rUTI Remission group membership. We then performed exploratory correlation analysis of creatinine-normalized estrogen metabolite concentrations and the species-level taxonomic profile. We found a striking difference in estrogen-associated taxonomic profiles between the No UTI History and rUTI Remission groups (Figure 5 E, F). We observed strong correlations between urinary E1 and E2 conjugates and *L. crispatus, L. iners, and L. gasseri* in the No UTI History group, correlations which were not detected in the rUTI Remission group (Figure 5 E, F Figure S 5 E, F). *B. breve*, an *Actinobacterium* associated with colon health, exhibited the strongest positive correlation across estrogen conjugates in the No UTI History group (Figure 5 E, G). This correlation was also absent in the urobiomes of the rUTI Remission group. *A. prevotii* was consistently and significantly negatively associated with urinary estrogens in the No UTI History group (Figure 5 E, Figure S 5 E). Fewer and distinct taxa correlated with estrogen conjugates in the rUTI Remission group (Figure 5 F, Figure S 5F). It should be noted that most EHT(+) women in the rUTI Remission group used vEHT (Figure 5 A). To assess the robustness of these results, we used an independent, Bayesian correlation approach to estimate normalized taxonomic abundances from read count-level data and account for overdispersion (Tu, 2014). Consistent with the non-Bayesian analysis, *L. gasseri, B. dentium, L. crispatus*, and *P. disiens* abundance were positively correlated with urinary estrogen conjugate sum in the No UTI history group, while *A. prevotii* was negatively correlated. Bayesian analysis also detected a set of new urinary estrogen-taxa correlations. Of note, *E. coli* was significantly anti-correlated with summed urinary estrogen concentration in the No UTI History group and *R. torques, Pantoea spp*., *Pseudomonas spp*., *Dorea spp*., and *C. aerofaciens* correlated with estrogen in the rUTI Remission group (Figure 5 H). Conversely, *D. microaerophilus* and *C. aurimucosum* were anti-correlated with urinary estrogen in the rUTI Remission group (Figure 5 I). Together, these data indicate that distinct urinary taxa correlate with urinary estrogen metabolites in women with No UTI history compared to women with rUTI history.

### Functional profiling reveals significant differences in the metabolic potential of cohort urobiomes

We next sought to determine if rUTI leaves a detectable imprint on the functional metabolic potential of the urobiome. We used the HUMAnN2 (v2.0) pipeline to profile the metabolic potential encoded within cohort urobiomes (Franzosa et al., 2018). Principal component analysis (PCA) performed on the relative abundance of encoded metabolic pathways in the three groups identified discriminating clusters that separated the rUTI Relapse urobiomes from the rUTI Remission and No UTI History urobiomes in the first two principal components (PCs) (Figure 6 A). These results were consistent with the taxonomic beta-diversity analysis (Figure 2 E). The rUTI Relapse group ordinated along vectors defined by the enrichment of lipopolysaccharide (LPS) biosynthesis (*n*=4 pathways), demethylmenaquinol-8 biosynthesis, fucose and mannose degradation, D-galacturonate degradation, sucrose degradation, and the TCA cycle (Figure 6 B). The rUTI Remission and No UTI History groups, which were not discriminated in the first two PCs, ordinated along vectors defined by the enrichment of nucleotide biosynthesis (*n*=8 pathways), L-lysine biosynthesis II, S-adenosyl methionine (SAM) biosynthesis, and UDP-N-acetyl-glucosamine biosynthesis (Figure 6 B). These data suggest that the large-scale genetic potential of the urobiome is relatively similar between rUTI Remission and No UTI History groups but is dramatically altered during active rUTI.

**Figure 6.**
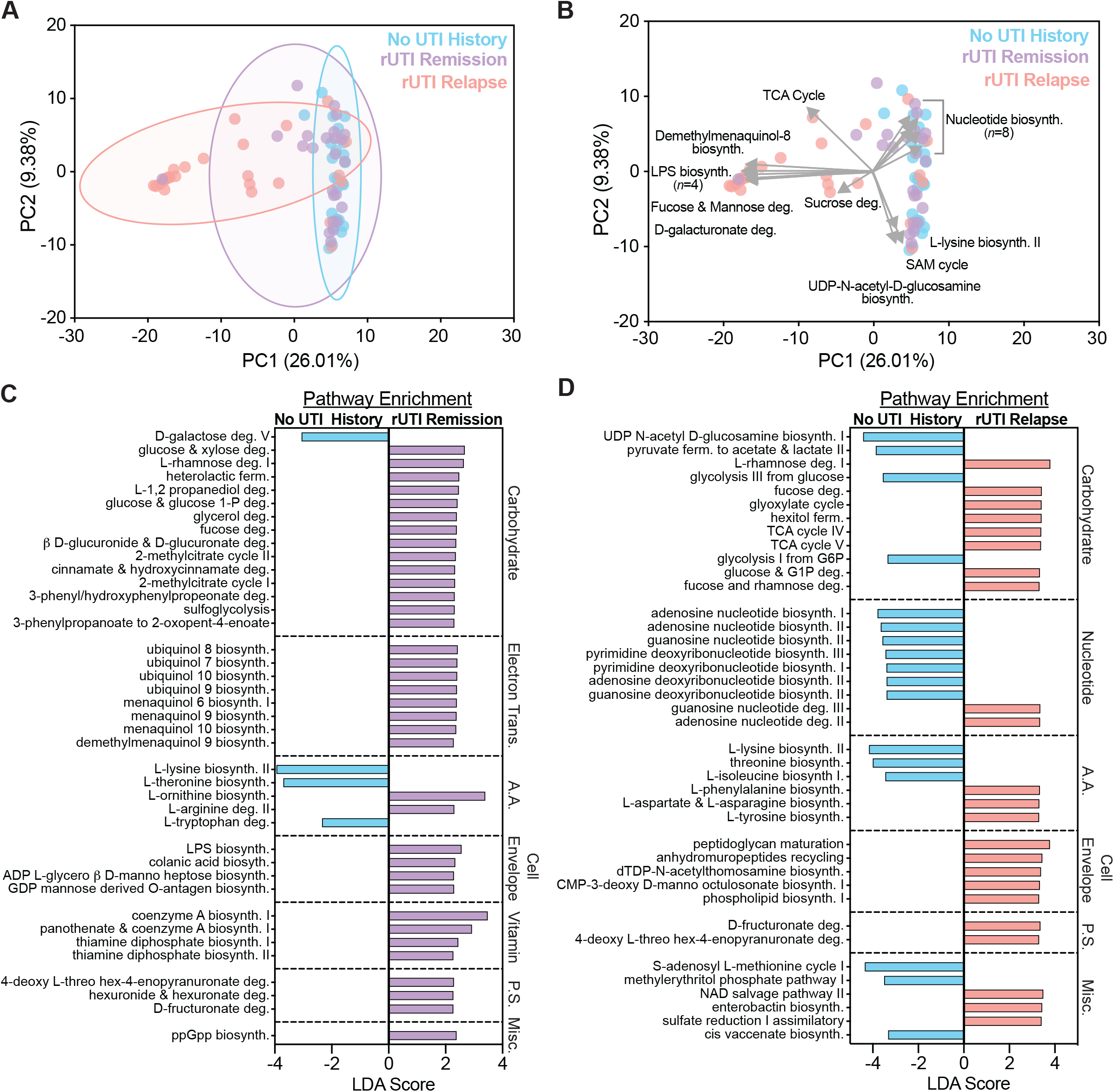
rUTI history and active infection shape the metabolic potential of the urobiome. (A, B) PCA of cohort urobiomes by metagenome-encoded metabolic pathways. Depiction of ordination and clustering in the first two PCs in (A) and vectors (grey) defining major discriminatory loadings (*i*.*e*. metabolic pathways) in (B). Top 40 differentially enriched pathways between No UTI history (blue) and rUTI Remission (purple) groups (C) and the No UTI history and rUTI Relapse (red) groups (D) detected by LEfSe. Pathways met an FDR-corrected *P*-value cutoff of < 0.05. LDA indicates log_10_(linear discriminant analysis).

Because dimensional reduction techniques often miss fine-scale discriminating features, we next used LEfSe to identify metabolic pathway enrichments between the study groups (Segata et al., 2011). First, we tested the hypothesis that rUTI history imparts functional changes on the urobiomes of PM women by comparing the No UTI History and rUTI Remission groups. Forty-five discriminatory metabolic pathways were significantly enriched in the rUTI Remission urobiomes and four metabolic pathways significantly enriched in the No UTI History urobiomes with an FDR-corrected *P*<0.05 and LDA >2 (Figure 6 C). The top 40 discriminating pathways were carbohydrate metabolism (*n*=14), electron carrier biosynthesis (*n*=8), amino acid metabolism (*n*=5), cell envelope biosynthesis (*n*=4), vitamin and cofactor biosynthesis (*n*=4), and polysaccharide degradation (*n*=3) (Figure 6 C). While most of the discriminatory carbohydrate metabolic pathways were enriched in the rUTI Remission urobiomes (13/14, 92.9%), we observed an enrichment of D-galactose degradation (Leloir pathway) in the No UTI History group (LDA=3.06, *P*=0.028). Conversely, electron carrier biosynthesis, namely biosynthetic pathways for ubiquinol 7-10 as well as menaquinol 6, 9 and 10 and demethylmenaquinol 9, were strongly enriched in the rUTI Remission urobiomes (Figure 6 C). L-lysine biosynthesis, L-threonine biosynthesis, and L-tryptophan degradation were enriched in No UTI History urobiomes, while L-ornithine biosynthesis and L-arginine degradation were enriched in rUTI Remission urobiomes. The remaining discriminating metabolic pathways were enriched in the rUTI Remission urobiomes and included cell envelope biosynthesis (e.g. LPS biosynthesis), vitamin metabolism (e.g. coenzyme A biosynthesis), polysaccharide degradation (e.g. 4-deoxy L-threo hex-4-enopyranuronate degradation), cinnamate and hydroxy cinnamate degradation, and ppGpp biosynthesis (Figure 6 C). These data suggest that the metabolic landscape of the urobiome may be significantly altered by rUTI history.

Pairwise differential enrichment analysis between the No UTI History and rUTI Relapse groups identified 183 differentially enriched metabolic pathways (Figure 6 D). In line with the taxonomic enrichment of Gram-negative species, we observed a strong enrichment of biosynthetic pathways for lipopolysaccharide (LPS) within rUTI Relapse urobiomes. Top discriminating pathways included carbohydrate (*n*=13), nucleotide (*n*=9), and amino acid metabolism (*n*=6), as well as cell envelope biosynthesis (*n*=5) (Figure 6 D). We observed enrichment of diverse carbohydrate degradation and central carbon metabolism pathways, including rhamnose, fucose, glyoxylate, and fructuronate degradation in rUTI Relapse urobiomes (Figure 6 D). This was coupled with a significant enrichment of TCA cycle metabolism, particularly 2-oxoglutarate decarboxylase and ferroreductase (Figure 6 D). Only four metabolic pathways involved in carbohydrate metabolism, including glycolysis from glucose and glucose 6-phosphate (G6P), pyruvate fermentation, and N-acetyl glucosamine biosynthesis, were significantly differentially enriched in the No UTI History urobiomes (Figure 6 D). Nucleic acid biosynthesis pathways were enriched in the No UTI History group while the Relapsed rUTI group was enriched for nucleic acid degradation pathways (Figure 6 D). Differentially enriched amino acid metabolism pathways included L-lysine, L-threonine, and L-isoleucine biosynthesis in the No UTI History group and L-phenylalanine biosynthesis in the rUTI Relapse group (Figure 6 D). These results suggest that the urobiomes of the rUTI Relapse group have the potential to utilize a more diverse nutrient set than the urobiomes of women who do not experience UTI.

### Antibiotic resistance genes are enriched in the urobiomes of women with rUTI history

Antibiotic therapy is currently the most prescribed treatment for the management of rUTI. However, resistance to front-line antibiotics, such as TMP-SMX, fluoroquinolones, and nitrofurantoin, is becoming a significant barrier to the successful treatment of rUTI (Malik et al., 2018a). We used the GROOT resistome analysis pipeline to generate a detailed profile of the antimicrobial resistance genes (ARGs) encoded within each urobiome (Rowe and Winn, 2018). This analysis detected 55 high-confidence ARGs distributed among all three groups. We observed significantly more ARGs in the urobiomes of the rUTI Remission (*P*=0.0455) and rUTI Relapse (*P*=0.0302) groups compared to No UTI History urobiomes (Figure 7 A). Interestingly, there was no significant difference in ARG count between rUTI Relapse and rUTI Remission urobiomes. These data suggest that a history of rUTI leaves an imprint on the underlying resistome of the urinary microbiota in PM women even in the absence of active infection.

**Figure 7.**
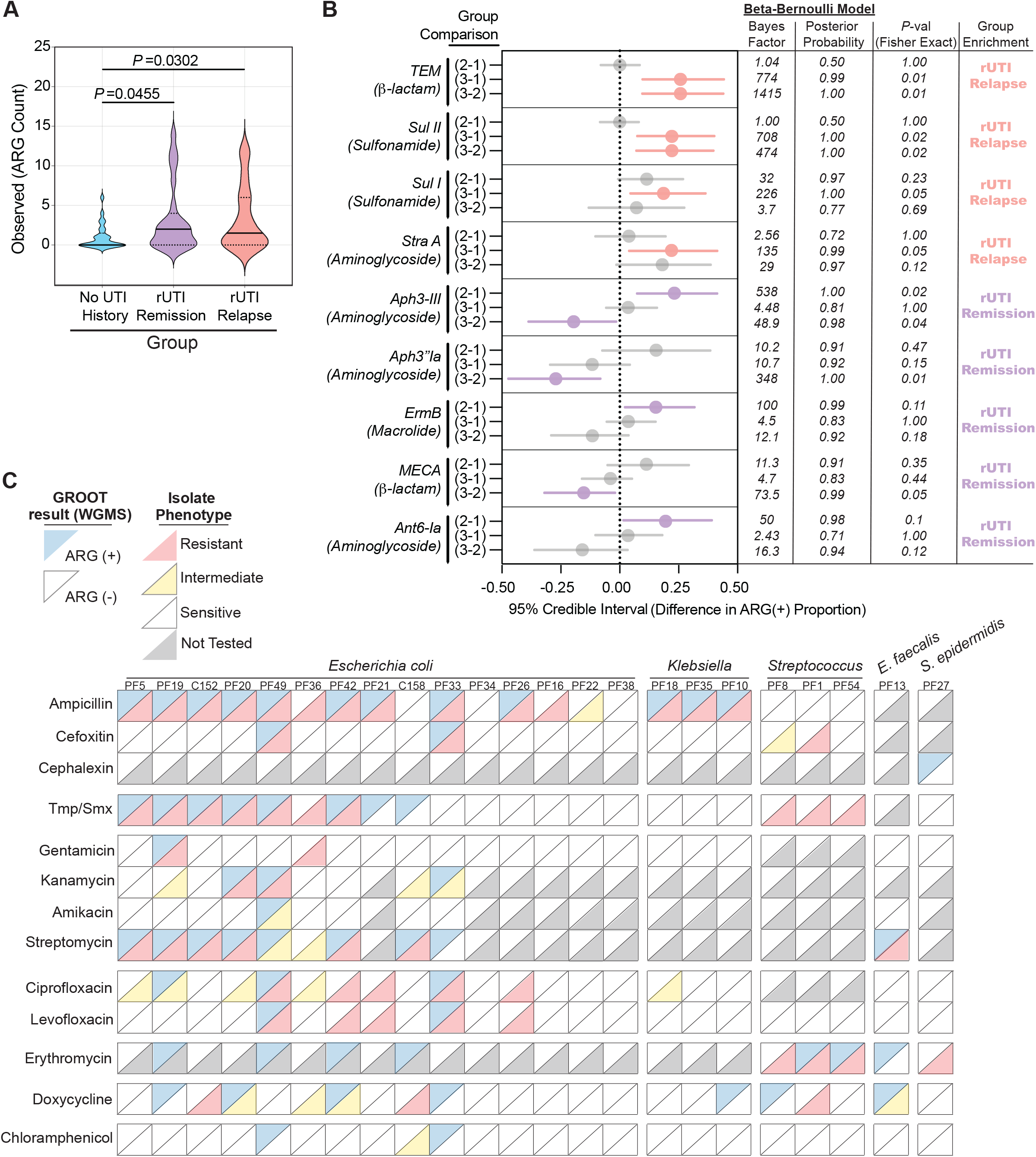
rUTI history and active infection shape the resistome of the PM urobiome. (A) Comparison of the ARGs detected within the urobiomes of the No UTI History, rUTI Remission, and rUTI Relapse groups. Violin plot depicts the smoothed distribution of the data. Solid lines represent median while dotted lines represent interquartile range. *P*-value generated by Kruskal-Wallis test with uncorrected Dunn’s multiple correction post hoc. (B) Bayesian differential enrichment analysis of ARGs within cohort urobiomes. Group comparisons were determined by pairwise differences in ARG(+) proportions. 95% credible intervals, Bayes factor, and posterior probability are presented for interpretation. Fisher exact *P*-values are also provided. (C) Agreement between urobiome ARG detection and antibiotic resistance phenotypes of isolates of the most abundant species present in each rUTI Relapse patient (*E. coli* (*n*=15), *Klebsiella* (*n*=3), *Streptococcus* (*n*=3), *E. faecali*s (*n*=1), *S. epidermidis* (*n*=1)). Upper diagonal colors represent WGMS profiling results (blue = ARG (+), white = ARG(-)). Lower diagonal color represents isolate phenotype (red = resistant, yellow = intermediate resistance, white = sensitive, grey = not tested).

To assess specific ARG enrichments associated with rUTI history, we performed differential enrichment analysis using a Bayesian model of proportional enrichment (Jeffreys, 1946). The TEM β-lactamase family, sulfonamide resistance genes *sul1* and *sul2*, and the *straA* aminoglycoside 3’-phosphotransferase were significantly enriched in the rUTI Relapse group while the aminoglycoside 3’-phosphotransferase genes, *aph(3’)-III* and *aph(3’)-Ia*, the macrolide resistance gene, *ermB*, the β-lactam resistance gene *mecA*, and the aminoglycoside O-nucleotidyltransferase gene, *ant(6)-Ia*, were enriched in the urobiomes of the rUTI Remission group (Figure 7 B). Conversely, no ARGs were significantly enriched in the No UTI History group. Identification of ARGs in metagenomes is only a prediction of the phenotype of a microbe or set of microbes belonging to the sampled population (Quince et al., 2017). To begin to understand how well metagenomic ARG analysis could predict phenotype, we measured antibiotic resistance phenotypes of 22 unique bacterial uropathogens each isolated from an individual rUTI Relapse patient. Species tested included *E. coli, K. pneumoniae, Klebsiella oxytoca, Streptococcus anginosus, S. agalactiae, E. faecalis*, and *S. epidermidis*. Only 3 of the 15 strains with complete or intermediate ampicillin resistance did not have a detected ampicillin resistance gene in the associated metagenome (Figure 7 C). WGMS resistome profiling detected cefixime resistance genes in 50% of the metagenomes (2/4) associated with isolates that were completely or intermediately resistant to cefixime (Figure 7 C). Interestingly, the cefixime resistant strains without a detected ARG were both streptococci. We observed that 50% of isolates exhibiting resistance to TMP/SMX were isolated from urobiomes for which WGMS resistome profiling detected the ARGs *sul I/II* and *drfA1* (Figure 7 C). In surveying aminoglycoside resistance, 50% (1/2), 60% (3/5), 100% (1/1), 88.9% (8/9) of the isolates exhibiting complete or intermediate resistance to Gentamicin, Kanamycin, Amikacin, and Streptomycin, respectively, were isolated from urobiomes with detected corresponding ARGs (Figure 7 C). Metagenomic ARG analysis was relatively poorly predictive for fluoroquinolone resistance with quinolone resistance genes detected in only 27.3% (3/11) and 33.3% (2/6) of the associated metagenomes of isolates with complete or intermediate resistance Ciprofloxacin and Levofloxacin, respectively (Figure 7 C). This is likely because GROOT does not detect single nucleotide polymorphisms (SNPs) and fluoroquinolone resistance is often conferred by SNPs in the genes encoding gyrase and topoisomerase I (Correia et al., 2017). Relatively few urobiomes harbored resistance factors to macrolides, which are not used in the treatment of Gram-negative infections (Arsic et al., 2018). While all Gram-positive bacterial strains were resistant to erythromycin, macrolide ARGs were only detected in the associated metagenomes of three strains. Tetracycline and phenicol ARGs were respectively detected in 42.9% (3/7) and 0% (0/1) of the metagenomes associated with strains with intermediate or resistant phenotypes (Figure 7C).

## Discussion

A decade of research has identified and characterized the urobiome (Brubaker and Wolfe, 2017; Hilt et al., 2014; Lewis et al., 2013; Price et al., 2019; Siddiqui et al., 2011; Wolfe et al., 2012). It has become evident that the urobiome is involved in the progression of, or affected by, urinary tract disease. Given the connection between host health and microbiome composition, the urobiome has drawn significant attention in further understanding rUTI susceptibility. Most WGMS studies of the female urobiome have been in younger or mixed age cohorts with little focused representation of PM women, leaving critical gaps in our knowledge of a heavily impacted demographic (Vaughan et al., 2021). Here, we present the first use of WGMS to specifically probe urobiome ecology and function associated with rUTI in PM women. We found that bacteria make up the majority of the non-human, non-viral metagenome of the PM urobiome and confirmed the viability of 93.9% of the genera observed at >5% relative abundance in the WGMS data by advanced urine culture. The main uropathogen detected in the rUTI Relapse group was UPEC, while urobiomes of the No UTI History and rUTI Remission groups were either dominated by a single bacterial species or were diverse. Both the No UTI History and rUTI Remission groups exhibited subsets of women with urobiomes dominated by *L. crispatus, L. gasseri, L. iners, B. breve, B. dentium, B. longum*, and *G. vaginalis*. These data support the observations by Thomas-White et al. of an interconnected urogenital microbiome (Thomas-White et al., 2018). We detected low abundances of fungal and archaeal taxa in all groups; however, little is known about the role of fungi and archaea in the female urobiome.

We identified differentially enriched urobiome taxa between healthy PM women and those with a history of rUTI that may serve as microbial biomarkers of urogenital tract health. Many of the genera enriched in the rUTI Remission group, including *Peptoniphilus, Prevotella, Actinomyces, Finegoldia, Fusobacterium*, and *Brevibacterium* were all members of the largest co-occurrence network that clustered strongly around the genus, *Peptoniphilus*, a known member of the vaginal microbiome associated with dysbiosis (Diop et al., 2019; Marrazzo et al., 2008; Onderdonk et al., 2016). These data suggest that mutualistic relationships between *Peptoniphilus* and co-occurring taxa may be key to defining community structure in urobiomes of women with increased rUTI susceptibility (Faust et al., 2012). Conversely, while the genus *Lactobacillus* and the species *L. vaginalis* and *L. crispatus* were enriched in No UTI History urobiomes, co-occurrence analysis revealed a negative association between *Lactobacillus* and *Peptoniphilus* suggesting an antagonistic relationship between the two genera. Taken together, these observations give new insight into potentially biologically relevant interactions in the urobiome that may underlie rUTI susceptibility. These observations are supported by a 2021 report by Vaughan et al. that used 16S rRNA amplicon sequencing to identify taxonomic differences between PM women with rUTI and healthy controls and identified differences in the orders *Clostridiales* and *Prevotellaeceae*, which, for example, contain the genera *Peptoniphilus* and *Prevotella*, respectively (Vaughan et al., 2021). Critical questions for future research will be to validate signatures of urinary dysbiosis associated with rUTI susceptibility in longitudinal studies.

Estrogen hormone therapy (EHT) is a common intervention to reduce discomfort associated with menopause (Fait, 2019). EHT, especially vEHT, is also gaining prevalence as a rUTI prophylactic in PM women because estrogen is thought to favor *Lactobacillus* colonization of the vaginal and urinary microbiomes (Raz and Stamm, 1993; Stamm, 2007). Multiple independent studies have evaluated associations between EHT and vaginal and urinary lactobacilli with varying results (Anglim et al., 2021; Thomas-White et al., 2020). For example, Anglim et al. found that vEHT did not significantly alter urinary lactobacilli populations among PM women with and without rUTI while Thomas-White et al. reported that vEHT led to a significant enrichment of urinary lactobacilli in PM women with overactive bladder symptoms (Anglim et al., 2021; Thomas-White et al., 2020). In the present study, we identify *L. crispatus and L. vaginalis* as uniquely associated with EHT use in PM women. Interestingly, streptococci as well as *A. vaginae*, a Gram-positive species associated with *G. vaginalis* in bacterial vaginosis, were enriched in the urobiomes of women not using EHT (Bradshaw et al., 2006). Importantly, careful analysis of clinical metadata found a strong link between oral and patch EHT use, but not vEHT use, and the presence of *Lactobacillus* in the urobiome. We hypothesized that EHT modalities may differ in dosage, composition, patient compliance, or primary metabolism. 17β-estradiol was the active ingredient in all oral and patch EHT formulations, but only 64.7% of vEHT(+) women used 17β-estradiol cream. The remaining vEHT(+) women used conjugated estrogen creams. Our data suggests that both oral and patch EHT are associated with elevated concentrations of urinary estrogen conjugates, while vEHT was not. We further identified disease state-specific taxa-estrogen metabolite correlations. *B. breve, L. iners, L. crispatus*, and *L. gasseri*, positively correlated with urinary estrogen conjugate concentration in the No UTI history group but not the rUTI Remission group. We hypothesize that this may be because rUTI Remission women were predominantly taking vEHT or because rUTI history itself perturbs the underlying urobiome. Future mechanistic research in relevant model systems and longitudinal human cohorts is needed to define the effects of modality of and rUTI history on EHT urobiome modulation.

Frequent and repeated treatment of rUTI with antibiotics is thought to spur the evolution of antibiotic resistance among uropathogenic bacteria and perhaps within the urobiome (Abbo and Hooton, 2014; Malik et al., 2018a; Waller et al., 2018). Despite the urgent need to understand the impact of antibiotic therapy on the urobiome, differences in urobiome ARG prevalence associated with rUTI and rUTI history had not been previously investigated. We found that the urobiomes of both the rUTI Relapse and rUTI Remission groups contained significantly more ARGs than the No UTI History group suggesting that rUTI history may enrich for ARG-containing urobiomes. Of note, rUTI Relapse urobiomes were enriched for β-lactam, aminoglycoside, and sulfonamide resistance genes while rUTI Remission urobiomes were enriched for β-lactam, aminoglycoside, macrolide resistance genes. Limitations of this analysis include that the analytical pipeline does not distinguish between TEM alleles and does not detect common SNPs known to confer fluoroquinolone resistance, for example.

Altogether, this work represents the first controlled WGMS analysis of urobiome structure and function in PM women with different histories of rUTI and provides a robust foundation for further mechanistic studies of the role of the urobiome in rUTI susceptibility and disease progression that are necessary for the development of urobiome-aware alternative therapies for a rUTI.

## Methods

### Patient Recruitment and Cohort Curation

The current study is approved under IRBs STU032016-006 (University of Texas Southwestern Medical Center) and 19MR0011 (University of Texas at Dallas). To estimate the number of patients needed to enroll into each cohort group and predict statistical power, we performed a series of power analyses using the ‘pwr’ package (https://github.com/heliosdrm/pwr) in the R-statistical language based on ranging effect sizes from small to large for both multivariate analysis of 3 groups with equal sample size and pairwise-based comparisons of two groups. Balancing cost and clinical feasibility with predicted statistical power, we chose a sample size of 25 for each cohort group. Patients were recruited from the Urology Clinic at the University of Texas Southwestern Medical Center between April 2018 and October 2019. Written informed consent was obtained from each patient prior to recruitment into the study cohorts. All patients were PM females. The following set of exclusion criteria were used to initially screen patient’s candidacy for enrollment into the cohort: pre- or perimenopausal status; complicated rUTI; antibiotic exposure within the 4 weeks prior to urine sample donation unless an active infection was detected by culture; pelvic malignancy or history of pelvic radiation within 3 years before urine sample donation; currently receiving chemotherapy; exhibiting renal insufficiency (creatinine >1.5 mg/dL); most recent post void residual (PVR) greater than 100 mL; greater than stage 2 prolapse; pelvic procedure for incontinence within 6 months prior to urine sample donation; use of intermittent catheterization; neurogenic bladder; any upper urinary tract abnormality which may explain rUTI; and Diabetes Mellitus (DM) type 1 or 2. All urine samples were obtained by clean-catch midstream urine collection and therefore were representative of the urogenital microbiome, rather than specifically just the bladder microbiome. Patients were educated about the cleaning and urine collection needs for this sampling technique prior to urine collection. Urine samples were stored at 4°C for no more than 4 hours before sample processing, aliquoting, and biobanking at -80°C. In total, 258 patients were recruited and screened for enrollment candidacy through interview, clinical assessment, and electronic medical records

Patients were further vetted by self-reporting of UTI history, mining clinical history from electronic patient records, clinical (standard) urine culture, and urine culture on Chromogenic agar (BBL CHROMagar Orientation, BD). Clinical urine culture was performed on samples from all patients with active UTI symptoms by the Clinical Microbiology Laboratory at UT Southwestern Medical Center. Group assignment criteria were as follows. No UTI History: no self-reported or clinical history of UTI, no UTI symptoms at the time of urine collection. rUTI Remission: recent history of rUTI, no UTI symptoms at time of sample collection. rUTI Relapse: recent history of rUTI, active UTI symptoms at time of urine collection, positive clinical urine culture. Positive urine culture was defined as >10^4^ bacterial CFU/mL.

### Advanced Urine Culture, Isolate Identification, and Isolate Biobanking

Glycerol-stocked urine samples (stored at -80ºC) were thawed at room temperature, and then diluted 1:3 and 1:30 in sterile 1X Phosphate Buffered Saline to adjust plating density for high and low biomass samples. 100 μl of urine from each dilution was plated onto blood agar plates (BAP), CHROMagar Orientation, De Man, Rogosa, and Sharpe (MRS) agar, Rabbit BAP (R-BAP), BD BBL CDC anaerobe blood agar (CDC AN-BAP), and Columbia Colistin Naladixic Acid Agar (CNA). Following plating, BAP was incubated in ambient and 5% CO_2_ atmospheres, CHROMagar Orientation in 5% CO_2_, MRS and R-BAP in microaerophilic conditions, BD BBL CDC anaerobe blood agar (CDC AN-BAP) in microaerophilic and anaerobic conditions and CNA in all four atmospheric conditions. Plates were incubated at 35°C for 4 days in the respective atmosphere. It should be noted that we were unable to culture *Gardnerella spp*. using these methods. However, WGMS profiling frequently detected *G. vaginalis* in the sampled urobiomes. For targeted isolation of *Gardnerella spp*.,100 μl urine was plated onto Human polysorbate-80 (HBT) bilayer medium in microaerophilic atmosphere for 3 days. To isolate fungal species, 100 μl urine was plated onto Brain Heart Infusion Agar supplemented with 20 g/L glucose and 50 mg/μl of chloramphenicol (BHIg-Cam) and incubated at 5% CO2 for 3 days.

Bacterial identification was performed by PCR amplification and Sanger sequencing of the 16S rRNA gene from well-isolated colonies as described previously (De Nisco et al., 2019). Briefly, 16S rRNA gene was amplified using primers 8F (5’-AGAGTTTGATCCTGGCTCAG-3’) and 1492R (5’-GGTTACCTTGTTACGACTT-3’) by colony PCR (Vaishnava et al., 2011) using DreamTaq Master Mix (ThermoFisher Scientific) and 0.2μM primers. Amplicon size was confirmed on 1% agarose gel, followed by gel purification (Bio basic) and Sanger Sequencing (Genewiz) using the 8F primer. Sequences were analyzed using BLASTn against the NCBI 16S ribosomal RNA (Bacteria and Archaea) database.

For fungal identification, ITS1 and ITS2 regions were amplified using the primer sequences ITS1: 5’-TCCGTAGGTGAACCTGCGG-3’ and ITS2: 5’-GCTGCGTTCTTCATCGATGC-3’ from well-isolated colonies and Sanger sequenced (Genewiz, South Plainfield, NJ, USA). Sequences were analyzed using BLASTn against the NCBI ITS from Fungi type and reference material database.

All the isolated and taxonomically identified isolates (*n*=896) were assigned a distinct ID and biobanked at -80ºC in glycerol. The isolates were grown in Brain Heart Infusion broth, Tryptic Soy Broth (BD 211825), MRS broth or NYCIII according to their growth preferences and stocked in 16% sterile glycerol for long-term storage at -80C.

### Metagenomic DNA Isolation, Library Construction, and Sequencing

Prior to WGMS, we assessed the quality and reproducibility of 3 metagenomic DNA extraction techniques: a modified genomic DNA (gDNA) isolation based on the Qiagen blood and tissue DNAeasy Kit, the Zymo Research DNA/RNA microbiome miniprep, and a modified phenol/chloroform/isoamyl alcohol extraction as demonstrated by Moustafa et al. (Moustafa et al., 2018). After assessing the quality and yield of metagenomic DNA isolated using the three methods, we chose the Zymo Research method. Urine samples were allowed to thaw on ice at 4°C. 10-20 mL of urine was centrifuged for 15 minutes at 4000 x g at 4°C. Urine pellets were resuspended in 750 μL of DNA/RNA Shield (Zymo Research), transferred to a bead beating tube, and subjected to ten 30 sec cycles of mechanical bead beating, with 5 min cooling between each cycle. After mechanical lysis, the maximum volume of sample was collected and transferred to a new microcentrifuge tube with DNA/RNA lysis buffer (Zymo). Nucleic acids were purified via the Zymo Research DNA/RNA microbiome miniprep kit per the manufacturer’s instruction. Elution of DNA from the column was performed in nuclease-free water and each column was eluted twice to maximize DNA recovery. As a control to internally assess gDNA extraction efficiency and WGMS limit of detection (LOD), gDNA was concurrently extracted from commercially available community standards (ZymoBiomics) using the same methods. gDNA was also extracted from nuclease-free water to account for kit and environmental contamination. All DNA samples were subjected to 16S rRNA gene amplification by PCR and visualized by agarose gel electrophoresis to ensure microbial DNA was present before proceeding with WGMS. DNA yield and purity for all samples were assessed by agarose gel electrophoresis, and by fluorescence-based Qubit quantitation of DNA, RNA, and protein. Prior to library preparation the DNA concentration of each sample was normalized and 20pg of spike-in gDNA was added (Zymo Research High Bacterial Load Spike-in), which contains gDNA from the bacterial species *Imchella halotolerans* and *Allobacillus halotolerans*, which are known to not be associated with humans.

WGMS was performed at the University of Texas at Dallas Genome Center using 2×150 bp paired-end reads on a Illumina NextSeq 500. Library preparation was performed using Nextera DNA Flex kit. Library preparation of the entire cohort and community standard and water controls was distributed over 2 batches with overlapping samples. All samples were sequenced using 2×150 base pair paired-end sequencing in high output mode with a target of ≥50 million paired end reads per sample.

### Bioinformatic Analyses

All taxonomic, functional, and resistome bioinformatic analyses were performed on an in-house Dell PowerEdge T630 server tower with 256GB RAM, 12 core Intel Xenon processor with 16TB storage capacity or at the Texas Advanced Computing Center (TACC).

#### Data Preprocessing

The fastq files were checked for read quality, adapter content, GC contents, species contamination using fastqc (v0.11.2) and fastq_screen (v0.4.4) (Andrews, 2015; Wingett and Andrews, 2018). Low-quality reads (a quality score of less than Q20) and adapter were removed using Trim galore (v 0.4.4) (Krueger) (Figure S 1D). Human DNA sequences were removed using KneadData (Huttenhower). After host removal, the dataset contained an average of 2.6×10^7^ non-human reads per sample.

#### Taxonomic Profiling, Ecological Modeling, and Co-Occurrence Analysis

The taxonomic assignment and estimation of composition of microbial species present in each sample was performed using MetaPhlAn2 (Segata et al., 2012). MetaPhlAn2 estimates the relative abundance of species by mapping the metagenomic reads against a clade specific marker gene database. The database consists of bacterial, archaeal, viral and eukaryotic genomes. We further used merge_metaphlan_tables module of MetaPhlAn2 to combine the relative abundance estimates of samples in a cohort into one table.

To identify kit, environmental, and background contaminating taxonomic signals, we sequenced a water sample which was randomly inserted into the metagenomic DNA preparation protocol. Sequencing and taxonomic analysis of this sample revealed known kit and environmental contaminants, such as *Delftia, Stenotrophomonas, Ralstonia, Bradyrizobium*, and others (Salter et al., 2014). Unless a known member of the human microbiome, these taxa were censored. We observed a small relative abundance of taxa salient to the human urobiome in the water control, such as *Pseudomonas, Escherichia, Klebsiella, Enterococcus, Staphylococcus*, and *Corynebacterium*. These signals ranged from 0.051%-11.5% of approximately 3 million mappable reads observed in the water control (Figure S 2A). We further assessed the WGMS limit of reliable detection using a commercially available log community standard (ZymoBiomics), which is composed of multiple Gram-positive and Gram-negative bacterial and fungi. We observed a strong linear correlation between the theoretical and observed relative abundance above 0.001%. We therefore set a relative abundance threshold of 0.001% for a taxon to be considered as detected within a sample (Figure S 1E). Species-level MetaPhlAn 2 taxonomic assignments were not included in analysis if they were “unclassified”.

Alpha-diversity analysis was performed at the species-level using phyloseq (version 1.16.2) (McMurdie and Holmes, 2013). Beta-diversity analysis was performed using DPCoA on the species-level taxonomic relative abundance dataset using phyloseq (version 1.16.2)(McMurdie and Holmes, 2013). Taxonomic co-occurrence was performed with CCREPE pipeline using the Pearson correlation and compositionally corrected P-values (https://github.com/biobakery/biobakery/wiki/ccrepe#22-ccrepe-function). Network analysis of taxonomic co-occurrences was performed using CytoScape (Version 3.8.2) with edges defined by the correlation coefficients between taxa nodes in the default prefuse force directed layout.

#### Taxonomic Correlation with Urinary Estrogens

The Spearman correlation was calculated between a given taxa and urinary estrogen conjugate or conjugate sum using the associate function from the regclass R package. To account for the compositionality of the taxa when computing the correlations, the species-level taxonomic composition dataset was transformed using the centered log ratio (CLR) transformation using the clr function from the rgr R package. Nominal P-values were calculated by permutation. No multiple hypothesis correction was performed on Nominal P-values as we considered this an exploratory analysis (Althouse, 2016).

The Bayesian correlation analysis employed a posterior distribution with the Dirichlet-Multinomial (DM) mode for the full data likelihood and a non-informative uniform prior proportional to one (Tu, 2014). The DM models the non-transformed count data directly and estimates their normalized abundances while accounting for overdispersion in the species-level count data. We then computed the Spearman correlation between each of the estimated normalized abundances and urinary estrogen conjugate sum. For posterior inference, we computed the 95% credible intervals and posterior means for the correlation between each of the normalized abundances and urinary estrogen conjugate sum. The correlation of a taxa-estrogen pair was significant if zero was not contained in the credible interval. Further for the significant pairs, we calculated the ratio of the proportion of posterior samples with correlations greater than 0.3 to those less than or equal to 0.3. This ratio (Posterior Ratio) indicates correlation strength where a ratio higher than one indicates a more moderate or strong correlation and less than one indicates a weak correlation.

#### Functional Metabolic Profiling

Functional metabolic profiling was performed using HUMAnN 2.0 (Franzosa et al., 2018). HUMAnN2 uses a tiered approach to identify the functional profile of microbial communities. Firstly, it maps the sample reads to clade specific markers and creates a database of pangenomes for each sample. In the second tier, it performs the nucleotide level mapping of samples reads against pangenome database. Lastly, a translated search against Uniref90 is performed for unaligned reads in each sample (Suzek et al., 2015). The output result is the mapping of reads to gene sequences with known taxonomy. The reads are normalized to gene sequence length to give an estimate of per-organism and community total gene family abundance. Next, gene families were analyzed to reconstruct and quantify metabolic pathways using MetaCyc (Caspi et al., 2018). Different modules of HUMAnN such as humann2_join_table and humann2_renorm_table were used to merge the pathway abundance of all the samples in a cohort and normalize the abundance to counts per million (cpm) respectively. We filtered the results to only include pathways whose taxonomic range included bacteria. We further censored pathways which were specifically associated with a particular taxon due to database bias toward commonly isolated and studied species. PCA of functional pathways was performed on the pathway level relative abundance dataset using factroextra (https://cran.r-project.org/web/packages/factoextra/readme/README.html). Pathway differential abundance analysis was performed using LEfSe (Segata et al., 2011) on the pathway-level relative abundance dataset. LEfSe uses Kruskal Wallis and Wilcoxon tests to find the differential pathways between microbial communities. Finally, it uses LDA model to rank the pathways.

#### Resistome Profiling and ARG Enrichment

We used the GROOT (Graphing Resistance Out Of meTagenomes) to generate a profile of antimicrobial resistance genes within the urobiomes of the present study (Rowe and Winn, 2018). The default database ARG-ANNOT was used for alignment of the metagenomics reads. Subsequently GROOT report command was used to generate a profile of antibiotic resistance genes at a read coverage of 90%. Filtering of the GROOT results was performed to insure high confidence in ARG presence within the urobiomes. We used a cutoff of a sufficient amount of reads to generate 10x coverage of an ARG to qualify its detection within a urobiome. We further collapsed alleles of the β-lactamase genes TEM, CTX, OXA, OXY2, SHV, and *cfxA* as well as the aminoglycoside ARG Aac3-IIa and Aac3-IIe alleles into single gene-level features to account for multiple-mapping reads.

Bayesian modeling of the resistome data was performed as follows. Resistome data for the three cohorts (Never = 1, Remission = 2, Relapse = 3) consisted of 186 antimicrobial resistance genes (ARG) which were collapsed into family-level genes (*G* = 55). Each cell in the data set contained a binary indicator of no detection (0) or detection (1) of the resistance family-level gene within each patient sample such that *x*_*gik*_ *=* {0,1}, *g* = 1, ⋯, 55, *i* = 1, ⋯, 25, *k* = 1,2,3 indicates no detection or detection of resistance family-level gene *g* respectively for sample *i* in cohort *k*.

A Bayesian Beta-Bernoulli model with Jeffreys prior was used to model the posterior distributions of group proportions and pairwise differences for the *g* family-level genes. Two posterior inferences were performed. First, we removed any family-level genes that had no significant pairwise contrast of cohort proportions using 95% credible intervals as criteria. We determined that a significant family level-gene does not have zero contained in a 95% credible interval for at least one pairwise contrast. Second, we computed the posterior probability and Bayes Factor (BF) to make inferences on each pairwise contrast of cohort proportions of only the significant family-level genes. The posterior probability of a particular contrast was computed as the proportion of posterior samples satisfying that contrast. The BF computed for each contrast represented the odds of *H*_1_: “at least one cohort’s proportion (ω) for gene *g* is different” in favor of *H*_0_: *ω*_*g*1_ *=ω*_*g*2_ *= ω*_*g*3_.

#### Taxonomic Biomarker Analysis

We applied two methods of taxonomic differential abundance analysis employing the robust and widely used LEfSe pipeline as well as BMDA, a recently described Bayesian model of differential abundance (Li et al., 2019). LEfSe analysis was performed as previously described (Segata et al., 2011). For the BMDA model we first applied the quality control step (detailed in the supplement of Li et al., 2019) to the raw count data. We then fitted the BMDA model, which is a Bayesian hierarchical framework that uses a zero-inflated binomial model to model the raw count data and a Gaussian mixture model with feature selection to identify differentially abundant taxa. The BMDA can fully account for zero-inflation, over-dispersion, and varying sequencing depth. We chose weakly informative priors on all parameters of the model to avoid biased results. For model fitting and posterior inference, BMDA implements the Metropolis-Hastings algorithm within a Gibbs sampler. The marginal posterior probability of inclusion (PPI) was used to identify the set of discriminating taxa between the control and disease groups. Marginal PPI is the proportion of MCMC samples in which a taxon is selected to be discriminatory if it is greater than a pre-specified value. We chose a threshold such that the expected Bayesian false discovery rate (FDR) was less than 0.05.

### Antibiotic susceptibility testing

Assessment of antibiotic (abx) susceptibility was performed via the Kirby-Bauer disk diffusion susceptibility test (Hudzicki, 2009). Antibiotic disks were prepared by aliquoting 10uL of antibiotic stock (GEN 1mg/ml, AMP 1mg/ml, CIP 0.5mg/ml, LVX 0.5mg/ml, ERM 1.5mg/ml, CHL 3mg/ml, TMP/SMX 1.25/23.75mg/ml, NIT 30mg/ml, DOX 3mg/ml) onto the disk in a sterile petri dish and drying at room temperature in the dark. Vehicle control disks were prepared similarly using the diluents of each antibiotic. Strains were streaked from frozen glycerol stocks onto CHROMagar or Blood Agar (species dependent) and incubated overnight at 37ºC in ambient conditions or 35ºC in 5% CO2. Single, well-isolated colonies were inoculated into 3 mL Brain-Heart-Infusion broth and incubated at the respective atmospheric conditions for 16 – 18 hours. After incubation, cultures were normalized to 0.5 McFarland standard, washed, and resuspended in sterile 1X Phosphate-Buffered Saline (PBS). 150 μL of standardized culture were pipetted onto 150 mm Mueller-Hinton Agar plates and spread using sterile glass beads. Plates were dried in sterile conditions before abx-impregnated disks were placed on the surface of the agar. *E. coli* strain ATCC25922 was used for quality and vehicle controls. Sterile 1X PBS was plated as sterility control. Plates were incubated inverted per the recommendations of Clinical and Laboratory Standards Institute (CLSI) M100-ED30: 2020 Performance Standards for Antimicrobial Susceptibility Testing, 30^th^ Edition (https://clsi.org/standards/products/microbiology/documents/m100/). After incubation, antimicrobial susceptibility was evaluated by measurement of the zone of inhibition and using CLSI established zone diameter breakpoints.

### Liquid Chromatography Mass Spectrometry Measurement of Estrogen Metabolites

Direct measurement of urinary estrogen metabolites was performed via a modification of previously reported methods(van der Berg et al., 2020). Briefly, urine (500 μL) was diluted and spiked with 100 ng stable isotope-labeled internal standards of d3-Estrone 3-Glucuronide and d4-Estradiol 3-Sulfate. Diluted and spiked samples were loaded onto an equilibrated Phenomenex C18 cartridge for solid phase extraction to separate conjugated estrogens. Following aqueous methanolic extraction of estrogen conjugates and non-polar extraction of free estrogens with methanolic acetone, fractions were dried by vacuum centrifugation and prepared for LC-MS/MS analysis. Estrogen conjugates (sulfates and glucuronides) were directly assayed using a curated and optimized MRM library by LC-MS/MS.

High sensitivity quantitative LC-MS/MS was performed on a Waters Xevo TQ tandem quadrupole MS lined to an ACQUITY UPLC with a Selectra C8 RP column (100×2.1 mm 1.8μm, UCT). MRM libraries of estrogen conjugates have been curated to include both analytical and confirmatory transitions for each analyte at optimal retention times to maximize separation. Briefly, data analysis was performed by integrating the peak area of the analytical transition for each analyte. Peak areas were normalized to molecular class-matched internal spike-in standards and mapped to a standard curve to accurately estimate analyte concentration. Urine estrogen metabolite concentrations were then normalized to urinary creatinine, which was measured by colorimetric assay (Sigma).

### Statistical Analysis

Statistical analysis was performed using R statistical programing, GraphPad Prism 9, and Microsoft Excel. For hypothesis testing, non-parametric Mann-Whitney U-test was used for pairwise comparisons and the Kruskal-Wallis non-parametric ANOVA with multiple comparison post-hoc was used for non-paired and unmatched comparisons of 3 or more groups. Multiple comparison adjustment was performed using false discover rate (FDR) when appropriate. An alpha of 0.05 was used to control type I error.

## Acknowledgements

We wish to acknowledge and sincerely thank the patients who participated in this study. This work was supported by a research grant from The Welch Foundation, a research grant from the Foundation for Women’s Wellness, NIH grant 1R01DK131267-01, and The University of Texas at Dallas Startup funds to NJD, the Cecil H. and Ida Green Chair in Systems Biology Science to KLP, The Felicia and John Cain Distinguished Chair in Women’s Health to PZ. We would also like to thank all the members of the De Nisco and Palmer labs for their helpful and creative input throughout this study. We would further like to acknowledge and thank the Genome Center at the University of Texas at Dallas for their invaluable work and assistance in generating the metagenomic dataset used for this study.

## Author Contributions

Conceptualization, M.L.N., P.E.Z., K.L.P., N.J.D.; data curation, M.L.N., J.F., A.Kup., A.P.A.; formal analysis, M.L.N, A.Kumar., K.C.L., C.Z., Q.L., C.X., V.S., N.J.D.; funding acquisition, V.S., K.L.P, P.E.Z., N.J.D.; investigation, M.L.N., N.V.H., V.H.N., A.N.; methodology, M.L.N., A.Kumar., N.V.H., K.C.L., V.H.N., A.N., B.M.S., Q.L., C.X., V.S., P.E.Z., K.L.P., N.J.D.; project administration, M.L.N., P.E.Z., N.J.D.; resources, P.E.Z., K.L.P., N.J.D.; software, M.L.N., A.Kumar., K.C.L., C.Z., Q.L., C.X.; supervision, Q.L., C.X., V.S., P.E.Z., K.L.P., N.J.D; validation, M.L.N., N.V.H., K.C.L., C.Z.; visualization, M.L.N., V.H.N.; writing-original draft, M.L.N., A.K., N.V.H., K.C.L., Q.L., C.X., N.J.D.

## Supplemental materials

Table S1. Cohort clinical characteristics

Figure S1. Power analysis and metagenomic dataset characteristics.

Figure S2. WGMS environmental contaminants and patient-level urine culturing coverage

Figure S3. Taxonomic profiles of detected *Archea, Eukaryota*, and Vial species.

Figure S4. Ecological modeling indices among the cohort groups.

Figure S5. Urinary estrogen conjugate concentrations and taxonomic associations.

## References

Abbo, L.M., and Hooton, T.M. (2014). Antimicrobial Stewardship and Urinary Tract Infections. Antibiotics (Basel) 3, 174–192.

Alteri, C.J., Himpsl, S.D., and Mobley, H.L. (2015). Preferential use of central metabolism in vivo reveals a nutritional basis for polymicrobial infection. PLoS Pathog 11, e1004601.

Althouse, A.D. (2016). Adjust for Multiple Comparisons? It’s Not That Simple. Ann Thorac Surg 101, 1644–1645.

Ammitzboll, N., Bau, B.P.J., Bundgaard-Nielsen, C., Villadsen, A.B., Jensen, A.M., Leutscher, P.D.C., Glavind, K., Hagstrom, S., Arenholt, L.T.S., and Sorensen, S. (2021). Pre- and postmenopausal women have different core urinary microbiota. Sci Rep 11, 2212.

Andrews, S. (2015). FastQC: a quality control tool for high throughput sequence data.

Anglim, B., Phillips, C., Shynlova, O., and Alarab, M. (2021). The effect of local estrogen therapy on the urinary microbiome composition of postmenopausal women with and without recurrent urinary tract infections. Int Urogynecol J.

Arsic, B., Barber, J., Cikos, A., Mladenovic, M., Stankovic, N., and Novak, P. (2018). 16-membered macrolide antibiotics: a review. Int J Antimicrob Agents 51, 283–298.

Barraud, O., Ravry, C., Francois, B., Daix, T., Ploy, M.C., and Vignon, P. (2019). Shotgun metagenomics for microbiome and resistome detection in septic patients with urinary tract infections. Int J Antimicrob Agents.

Bradshaw, C.S., Tabrizi, S.N., Fairley, C.K., Morton, A.N., Rudland, E., and Garland, S.M. (2006). The association of Atopobium vaginae and Gardnerella vaginalis with bacterial vaginosis and recurrence after oral metronidazole therapy. J Infect Dis 194, 828–836.

Brubaker, L., and Wolfe, A.J. (2017). The female urinary microbiota, urinary health and common urinary disorders. Ann Transl Med 5, 34.

Bucevic Popovic, V., Situm, M., Chow, C.T., Chan, L.S., Roje, B., and Terzic, J. (2018). The urinary microbiome associated with bladder cancer. Sci Rep 8, 12157.

Caspi, R., Billington, R., Fulcher, C.A., Keseler, I.M., Kothari, A., Krummenacker, M., Latendresse, M., Midford, P.E., Ong, Q., Ong, W.K., et al. (2018). The MetaCyc database of metabolic pathways and enzymes. Nucleic Acids Res 46, D633–D639.

Ceccarani, C., Foschi, C., Parolin, C., D’Antuono, A., Gaspari, V., Consolandi, C., Laghi, L., Camboni, T., Vitali, B., Severgnini, M., et al. (2019). Diversity of vaginal microbiome and metabolome during genital infections. Sci Rep 9, 14095.

Correia, S., Poeta, P., Hebraud, M., Capelo, J.L., and Igrejas, G. (2017). Mechanisms of quinolone action and resistance: where do we stand? J Med Microbiol 66, 551–559.

De Nisco, N.J., Neugent, M., Mull, J., Chen, L., Kuprasertkul, A., de Souza Santos, M., Palmer, K.L., Zimmern, P., and Orth, K. (2019). Direct Detection of Tissue-Resident Bacteria and Chronic Inflammation in the Bladder Wall of Postmenopausal Women with Recurrent Urinary Tract Infection. J Mol Biol 431, 4368–4379.

Diop, K., Diop, A., Michelle, C., Richez, M., Rathored, J., Bretelle, F., Fournier, P.E., and Fenollar, F. (2019). Description of three new Peptoniphilus species cultured in the vaginal fluid of a woman diagnosed with bacterial vaginosis: Peptoniphilus pacaensis sp. nov., Peptoniphilus raoultii sp. nov., and Peptoniphilus vaginalis sp. nov. Microbiologyopen 8, e00661.

Edwards, V.L., Smith, S.B., McComb, E.J., Tamarelle, J., Ma, B., Humphrys, M.S., Gajer, P., Gwilliam, K., Schaefer, A.M., Lai, S.K., et al. (2019). The Cervicovaginal Microbiota-Host Interaction Modulates Chlamydia trachomatis Infection. MBio 10.

Fait, T. (2019). Menopause hormone therapy: latest developments and clinical practice. Drugs Context 8, 212551.

Faust, K., Sathirapongsasuti, J.F., Izard, J., Segata, N., Gevers, D., Raes, J., and Huttenhower, C. (2012). Microbial co-occurrence relationships in the human microbiome. PLoS Comput Biol 8, e1002606.

Flores-Mireles, A.L., Walker, J.N., Caparon, M., and Hultgren, S.J. (2015). Urinary tract infections: epidemiology, mechanisms of infection and treatment options. Nat Rev Microbiol 13, 269–284.

Franzosa, E.A., McIver, L.J., Rahnavard, G., Thompson, L.R., Schirmer, M., Weingart, G., Lipson, K.S., Knight, R., Caporaso, J.G., Segata, N., et al. (2018). Species-level functional profiling of metagenomes and metatranscriptomes. Nat Methods 15, 962–968.

Gaitonde, S., Malik, R.D., and Zimmern, P.E. (2019). Financial Burden of Recurrent Urinary Tract Infections in Women: A Time-driven Activity-based Cost Analysis. Urology 128, 47–54.

Hardy, L., Jespers, V., Abdellati, S., De Baetselier, I., Mwambarangwe, L., Musengamana, V., van de Wijgert, J., Vaneechoutte, M., and Crucitti, T. (2016). A fruitful alliance: the synergy between Atopobium vaginae and Gardnerella vaginalis in bacterial vaginosis-associated biofilm. Sex Transm Infect 92, 487–491.

Hardy, L., Jespers, V., Dahchour, N., Mwambarangwe, L., Musengamana, V., Vaneechoutte, M., and Crucitti, T. (2015). Unravelling the Bacterial Vaginosis-Associated Biofilm: A Multiplex Gardnerella vaginalis and Atopobium vaginae Fluorescence In Situ Hybridization Assay Using Peptide Nucleic Acid Probes. PLoS One 10, e0136658.

Hilt, E.E., McKinley, K., Pearce, M.M., Rosenfeld, A.B., Zilliox, M.J., Mueller, E.R., Brubaker, L., Gai, X., Wolfe, A.J., and Schreckenberger, P.C. (2014). Urine is not sterile: use of enhanced urine culture techniques to detect resident bacterial flora in the adult female bladder. J Clin Microbiol 52, 871–876.

Hudzicki, J. (2009). Kirby-Bauer disk diffusion susceptibility test protocol.

Huttenhower, C. KneadData.

Jeffreys, H. (1946). An invariant form for the prior probability in estimation problems. Proc R Soc Lond A Math Phys Sci 186, 453–461.

Jhang, J.F., and Kuo, H.C. (2017). Recent advances in recurrent urinary tract infection from pathogenesis and biomarkers to prevention. Ci Ji Yi Xue Za Zhi 29, 131–137.

Karstens, L., Asquith, M., Caruso, V., Rosenbaum, J.T., Fair, D.A., Braun, J., Gregory, W.T., Nardos, R., and McWeeney, S.K. (2018). Community profiling of the urinary microbiota: considerations for low-biomass samples. Nat Rev Urol 15, 735–749.

Karstens, L., Asquith, M., Davin, S., Stauffer, P., Fair, D., Gregory, W.T., Rosenbaum, J.T., McWeeney, S.K., and Nardos, R. (2016). Does the Urinary Microbiome Play a Role in Urgency Urinary Incontinence and Its Severity? Front Cell Infect Microbiol 6, 78.

Keogh, D., Tay, W.H., Ho, Y.Y., Dale, J.L., Chen, S., Umashankar, S., Williams, R.B.H., Chen, S.L., Dunny, G.M., and Kline, K.A. (2016). Enterococcal Metabolite Cues Facilitate Interspecies Niche Modulation and Polymicrobial Infection. Cell Host Microbe 20, 493–503.

Kononen, E., and Wade, W.G. (2015). Actinomyces and related organisms in human infections. Clin Microbiol Rev 28, 419–442.

Krueger, F. Trim Galore!

Lewis, D.A., Brown, R., Williams, J., White, P., Jacobson, S.K., Marchesi, J.R., and Drake, M.J. (2013). The human urinary microbiome; bacterial DNA in voided urine of asymptomatic adults. Front Cell Infect Microbiol 3, 41.

Li, Q., Jiang, S., Koh, A.Y., Xiao, G., and Zhan, X. (2019). Bayesian Modeling of Microbiome Data for Differential Abundance Analysis, pp. 1902.08741.

Lobo, R.A. (2017). Hormone-replacement therapy: current thinking. Nat Rev Endocrinol 13, 220–231.

Malik, R.D., Wu, Y.R., Christie, A.L., Alhalabi, F., and Zimmern, P.E. (2018a). Impact of Allergy and Resistance on Antibiotic Selection for Recurrent Urinary Tract Infections in Older Women. Urology 113, 26–33.

Malik, R.D., Wu, Y.R., and Zimmern, P.E. (2018b). Definition of Recurrent Urinary Tract Infections in Women: Which One to Adopt? Female Pelvic Med Reconstr Surg 24, 424–429.

Marrazzo, J.M., Thomas, K.K., Fiedler, T.L., Ringwood, K., and Fredricks, D.N. (2008). Relationship of specific vaginal bacteria and bacterial vaginosis treatment failure in women who have sex with women. Ann Intern Med 149, 20–28.

McMurdie, P.J., and Holmes, S. (2013). phyloseq: an R package for reproducible interactive analysis and graphics of microbiome census data. PLoS One 8, e61217.

Moustafa, A., Li, W., Singh, H., Moncera, K.J., Torralba, M.G., Yu, Y., Manuel, O., Biggs, W., Venter, J.C., Nelson, K.E., et al. (2018). Microbial metagenome of urinary tract infection. Sci Rep 8, 4333.

Neugent, M.L., Hulyalkar, N.V., Nguyen, V.H., Zimmern, P.E., and De Nisco, N.J. (2020). Advances in Understanding the Human Urinary Microbiome and Its Potential Role in Urinary Tract Infection. mBio 11.

Onderdonk, A.B., Delaney, M.L., and Fichorova, R.N. (2016). The Human Microbiome during Bacterial Vaginosis. Clin Microbiol Rev 29, 223–238.

Pavoine, S., Dufour, A.B., and Chessel, D. (2004). From dissimilarities among species to dissimilarities among communities: a double principal coordinate analysis. J Theor Biol 228, 523–537.

Pearce, M.M., Hilt, E.E., Rosenfeld, A.B., Zilliox, M.J., Thomas-White, K., Fok, C., Kliethermes, S., Schreckenberger, P.C., Brubaker, L., Gai, X., et al. (2014). The female urinary microbiome: a comparison of women with and without urgency urinary incontinence. MBio 5, e01283–01214.

Price, T.K., Dune, T., Hilt, E.E., Thomas-White, K.J., Kliethermes, S., Brincat, C., Brubaker, L., Wolfe, A.J., Mueller, E.R., and Schreckenberger, P.C. (2016). The Clinical Urine Culture: Enhanced Techniques Improve Detection of Clinically Relevant Microorganisms. J Clin Microbiol 54, 1216–1222.

Price, T.K., Hilt, E.E., Thomas-White, K., Mueller, E.R., Wolfe, A.J., and Brubaker, L. (2019). The urobiome of continent adult women: a cross-sectional study. BJOG.

Quince, C., Walker, A.W., Simpson, J.T., Loman, N.J., and Segata, N. (2017). Shotgun metagenomics, from sampling to analysis. Nat Biotechnol 35, 833–844.

Ravel, J., and Brotman, R.M. (2016). Translating the vaginal microbiome: gaps and challenges. Genome Med 8, 35.

Ravel, J., Gajer, P., Abdo, Z., Schneider, G.M., Koenig, S.S., McCulle, S.L., Karlebach, S., Gorle, R., Russell, J., Tacket, C.O., et al. (2011). Vaginal microbiome of reproductive-age women. Proc Natl Acad Sci U S A 108 Suppl 1, 4680–4687.

Raz, R., and Stamm, W.E. (1993). A controlled trial of intravaginal estriol in postmenopausal women with recurrent urinary tract infections. N Engl J Med 329, 753–756.

Rowe, W.P.M., and Winn, M.D. (2018). Indexed variation graphs for efficient and accurate resistome profiling. Bioinformatics 34, 3601–3608.

Salter, S.J., Cox, M.J., Turek, E.M., Calus, S.T., Cookson, W.O., Moffatt, M.F., Turner, P., Parkhill, J., Loman, N.J., and Walker, A.W. (2014). Reagent and laboratory contamination can critically impact sequence-based microbiome analyses. BMC Biol 12, 87.

Segata, N., Izard, J., Waldron, L., Gevers, D., Miropolsky, L., Garrett, W.S., and Huttenhower, C. (2011). Metagenomic biomarker discovery and explanation. Genome Biol 12, R60.

Segata, N., Waldron, L., Ballarini, A., Narasimhan, V., Jousson, O., and Huttenhower, C. (2012). Metagenomic microbial community profiling using unique clade-specific marker genes. Nat Methods 9, 811–814.

Shipitsyna, E., Roos, A., Datcu, R., Hallen, A., Fredlund, H., Jensen, J.S., Engstrand, L., and Unemo, M. (2013). Composition of the vaginal microbiota in women of reproductive age--sensitive and specific molecular diagnosis of bacterial vaginosis is possible? PLoS One 8, e60670.

Siddiqui, H., Nederbragt, A.J., Lagesen, K., Jeansson, S.L., and Jakobsen, K.S. (2011). Assessing diversity of the female urine microbiota by high throughput sequencing of 16S rDNA amplicons. BMC Microbiol 11, 244.

Stamm, W.E. (2007). Estrogens and urinary-tract infection. J Infect Dis 195, 623–624.

Stamm, W.E., and Norrby, S.R. (2001). Urinary tract infections: disease panorama and challenges. J Infect Dis 183 Suppl 1, S1–4.

Stapleton, A.E., Au-Yeung, M., Hooton, T.M., Fredricks, D.N., Roberts, P.L., Czaja, C.A., Yarova-Yarovaya, Y., Fiedler, T., Cox, M., and Stamm, W.E. (2011). Randomized, placebo-controlled phase 2 trial of a Lactobacillus crispatus probiotic given intravaginally for prevention of recurrent urinary tract infection. Clin Infect Dis 52, 1212–1217.

Suzek, B.E., Wang, Y., Huang, H., McGarvey, P.B., Wu, C.H., and UniProt, C. (2015). UniRef clusters: a comprehensive and scalable alternative for improving sequence similarity searches. Bioinformatics 31, 926–932.

Thomas-White, K., Forster, S.C., Kumar, N., Van Kuiken, M., Putonti, C., Stares, M.D., Hilt, E.E., Price, T.K., Wolfe, A.J., and Lawley, T.D. (2018). Culturing of female bladder bacteria reveals an interconnected urogenital microbiota. Nat Commun 9, 1557.

Thomas-White, K., Taege, S., Limeira, R., Brincat, C., Joyce, C., Hilt, E.E., Mac-Daniel, L., Radek, K.A., Brubaker, L., Mueller, E.R., et al. (2020). Vaginal estrogen therapy is associated with increased Lactobacillus in the urine of postmenopausal women with overactive bladder symptoms. Am J Obstet Gynecol 223, 727 e721–727 e711.

Thomas-White, K.J., Kliethermes, S., Rickey, L., Lukacz, E.S., Richter, H.E., Moalli, P., Zimmern, P., Norton, P., Kusek, J.W., Wolfe, A.J., et al. (2017). Evaluation of the urinary microbiota of women with uncomplicated stress urinary incontinence. Am J Obstet Gynecol 216, 55 e51–55 e16.

Tu, S. (2014). The Dirichlet-Multinomial and Dirichlet-Categorical models for Bayesian inference.

Vaishnava, S., Yamamoto, M., Severson, K.M., Ruhn, K.A., Yu, X., Koren, O., Ley, R., Wakeland, E.K., and Hooper, L.V. (2011). The antibacterial lectin RegIIIgamma promotes the spatial segregation of microbiota and host in the intestine. Science 334, 255–258.

van der Berg, C.L., Venter, G., van der Westhuizen, F.H., and Erasmus, E. (2020). Data on the optimisation of a solid phase extraction method for fractionating estrogen metabolites from small urine volumes. Data Brief 29, 105222.

Vaughan, M.H., Mao, J., Karstens, L.A., Ma, L., Amundsen, C.L., Schmader, K.E., and Siddiqui, N.Y. (2021). The Urinary Microbiome in Postmenopausal Women with Recurrent Urinary Tract Infections. J Urol 206, 1222–1231.

Waller, T.A., Pantin, S.A.L., Yenior, A.L., and Pujalte, G.G.A. (2018). Urinary Tract Infection Antibiotic Resistance in the United States. Prim Care 45, 455–466.

Wingett, S.W., and Andrews, S. (2018). FastQ Screen: A tool for multi-genome mapping and quality control. F1000Res 7, 1338.

Wolfe, A.J., and Brubaker, L. (2019). Urobiome updates: advances in urinary microbiome research. Nat Rev Urol 16, 73–74.

Wolfe, A.J., Toh, E., Shibata, N., Rong, R., Kenton, K., Fitzgerald, M., Mueller, E.R., Schreckenberger, P., Dong, Q., Nelson, D.E., et al. (2012). Evidence of uncultivated bacteria in the adult female bladder. J Clin Microbiol 50, 1376–1383.

