## Supplemental Figures and Legends for "Recurrent urinary tract infection and estrogen shape the taxonomic ecology and functional potential of the postmenopausal urobiome"

**Supplemental Table 1**

| | No UTI History (Group 1) | rUTI Remission (Group 2) | rUTI Relapse (Group 3) | *P*-value |
|---|---|---|---|---|
| ***N*** **(Individuals)** | 25 | 25 | 25 | **-** |
| **Age (years) ($CI_{95\%}$)** | 67 (62-73) | 68 (64-77) | 76 (70-78) | **0.04**[A] |
| **Race** | | | | |
| *African American* | 1 (4%) | 1 (4%) | 0 (0%) | |
| *Caucasian* | 24 (96%) | 23 (92%) | 22 (88%) | |
| *Hispanic* | 0 (0%) | 1 (4%) | 3 (12%) | |
| *Other* | 0 (0%) | 0 (0%) | 0 (0%) | 0.33[B] |
| **BMI ($CI_{95\%}$)** | 26.2 (23.7-28.9) | 25.3 (23.0-29.1) | 27.3 (23.0-29.1) | 0.25[A] |
| **Smoking History** | | | | |
| *Never* | 16 (64%) | 17 (68%) | 17 (68%) | |
| *Ever* | 9 (36%) | 8 (32%) | 8 (32%) | 0.94[B] |
| **EHT** | | | | |
| *EHT (-)* | 10 (40%) | 11 (44%) | 17 (68%) | |
| *EHT (+)* | 15 (60%) | 14 (56%) | 8 (32%) | 0.10[B] |
| **Urine pH ($CI_{95\%}$)** | 6.0 (5.0-7.0) | 6.0 (5.0-6.9) | 5.5 (5.0-6.3) | 0.60[A] |
| **Urinary Creatinine (μg/ml) ($CI_{95\%}$)** | 644.9 (301.2-1007) | 720.7 (419.8-957.1) | 771.5 (483.0-1083) | 0.69[A] |

[A] Kruskal-Wallis test
[B] $\chi^2$ test

### Figure S1

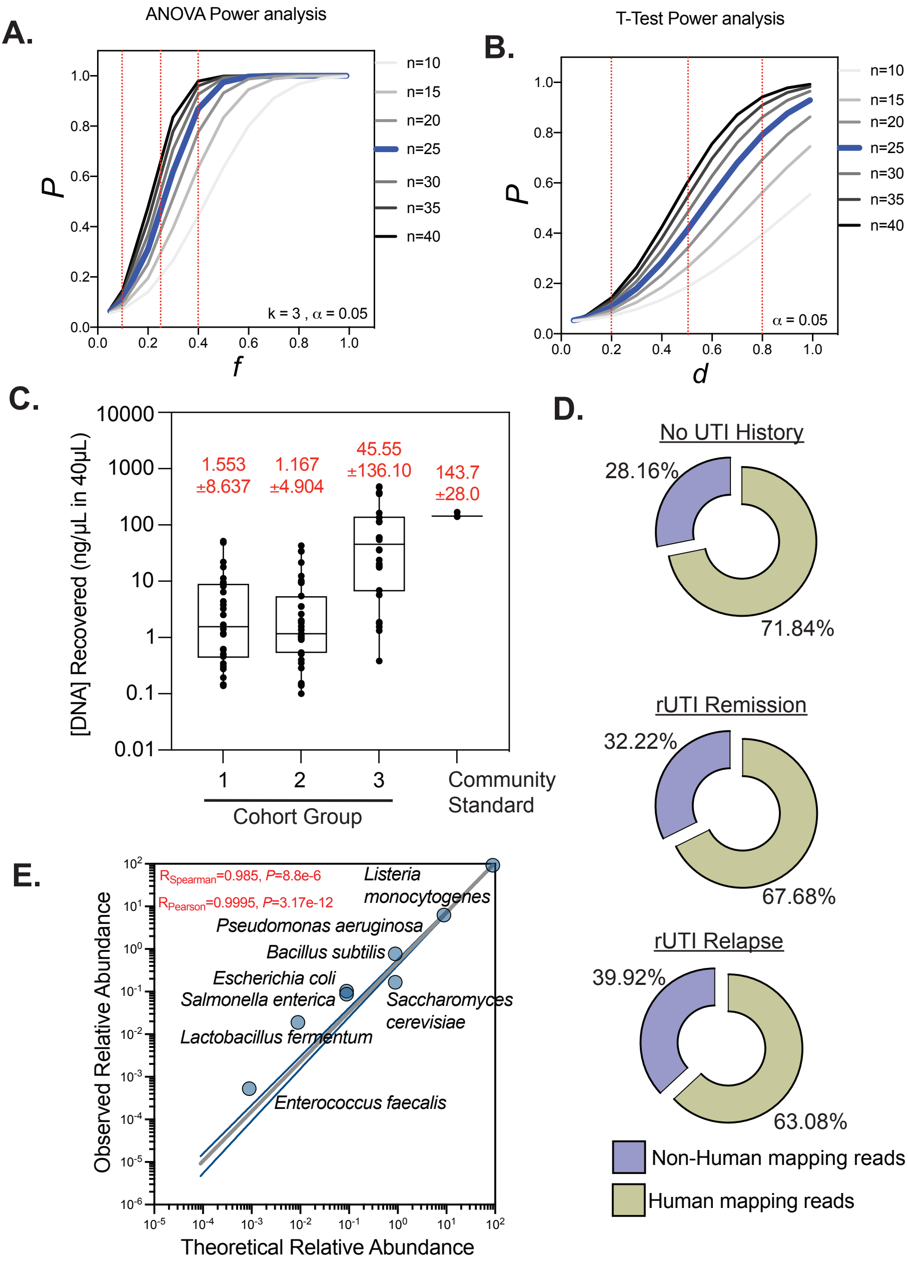

A.
ANOVA Power analysis
P
f
k = 3 , α = 0.05
n=10
n=15
n=20
n=25
n=30
n=35
n=40
B.
T-Test Power analysis
P
d
α = 0.05
n=10
n=15
n=20
n=25
n=30
n=35
n=40
C.
[DNA] Recovered (ng/μL in 40μL)
1.553 ±8.637
1.167 ±4.904
45.55 ±136.10
143.7 ±28.0
1
2
3
Community Standard
Cohort Group
D.
No UTI History
28.16%
71.84%
rUTI Remission
32.22%
67.68%
rUTI Relapse
39.92%
63.08%
Non-Human mapping reads
Human mapping reads
E.
R_Spearman=0.985, P=8.8e-6
R_Pearson=0.9995, P=3.17e-12
Listeria monocytogenes
Pseudomonas aeruginosa
Bacillus subtilis
Escherichia coli
Salmonella enterica
Saccharomyces cerevisiae
Lactobacillus fermentum
Enterococcus faecalis
Observed Relative Abundance
Theoretical Relative Abundance

**Figure S2**

A

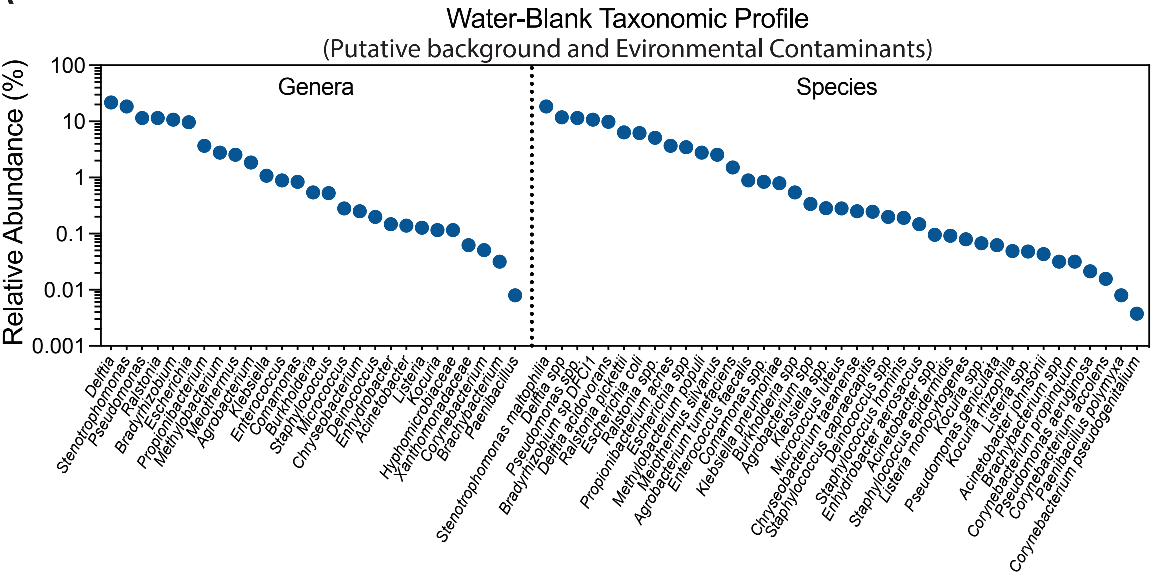

Water-Blank Taxonomic Profile
(Putative background and Evironmental Contaminants)
Relative Abundance (%)
100
10
1
0.1
0.01
0.001
Genera
Species
Delftia
Stenotrophomonas
Pseudomonas
Ralstonia
Bradyrhizobium
Escherichia
Propionibacterium
Methylobacterium
Meiothermus
Agrobacterium
Klebsiella
Enterococcus
Comamonas
Burkholderia
Staphylococcus
Micrococcus
Chryseobacterium
Deinococcus
Enhydrobacter
Acinetobacter
Listeria
Kocuria
Hyphomicrobiaceae
Xanthomonadaceae
Corynebacterium
Brachybacterium
Paenibacillus
Stenotrophomonas maltophilia
Delftia spp
Pseudomonas spp.
Bradyrhizobium sp DFCI1
Delftia acidovorans
Ralstonia pickettii
Escherichia coli
Ralstonia spp.
Propionibacterium acnes
Escherichia spp
Methylobacterium populi
Meiothermus silvanus
Agrobacterium tumefaciens
Enterococcus faecalis
Comamonas spp.
Klebsiella pneumoniae
Burkholderia spp
Agrobacterium spp
Klebsiella spp.
Micrococcus luteus
Chryseobacterium taeanense
Staphylococcus capraecapitis
Deinococcus spp
Staphylococcus hominis
Enhydrobacter aerosaccus
Acinetobacter spp.
Staphylococcus epidermidis
Listeria monocytogenes
Kocuria spp.
Pseudomonas geniculata
Kocuria rhizophila
Listeria spp.
Acinetobacter johnsonii
Brachybacterium spp
Corynebacterium propinquum
Pseudomonas aeruginosa
Corynebacterium accolens
Paenibacillus polymyxa
Corynebacterium pseudogenitalium


B

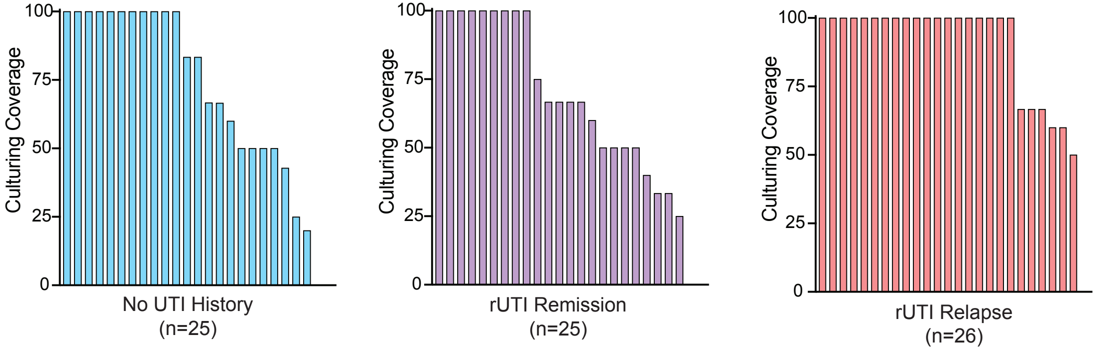

Culturing Coverage
100
75
50
25
0
No UTI History
(n=25)
rUTI Remission
(n=25)
rUTI Relapse
(n=26)

### Figure S3

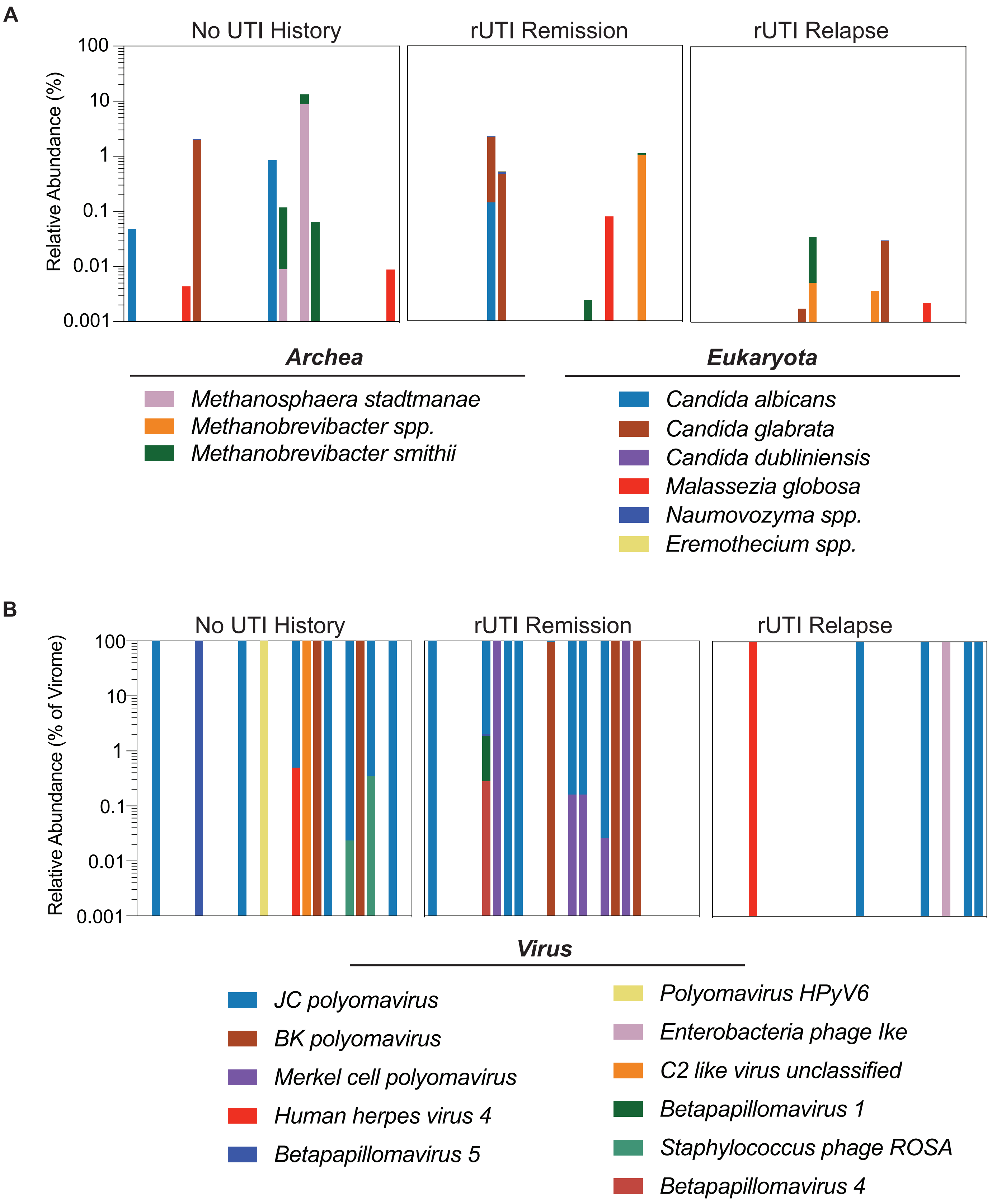
A
No UTI History
rUTI Remission
rUTI Relapse
Relative Abundance (%)
100
10
1
0.1
0.01
0.001
Archea
Methanosphaera stadtmanae
Methanobrevibacter spp.
Methanobrevibacter smithii
Eukaryota
Candida albicans
Candida glabrata
Candida dubliniensis
Malassezia globosa
Naumovozyma spp.
Eremothecium spp.
B
No UTI History
rUTI Remission
rUTI Relapse
Relative Abundance (% of Virome)
100
10
1
0.1
0.01
0.001
Virus
JC polyomavirus
BK polyomavirus
Merkel cell polyomavirus
Human herpes virus 4
Betapapillomavirus 5
Polyomavirus HPyV6
Enterobacteria phage Ike
C2 like virus unclassified
Betapapillomavirus 1
Staphylococcus phage ROSA
Betapapillomavirus 4

**Figure S4**

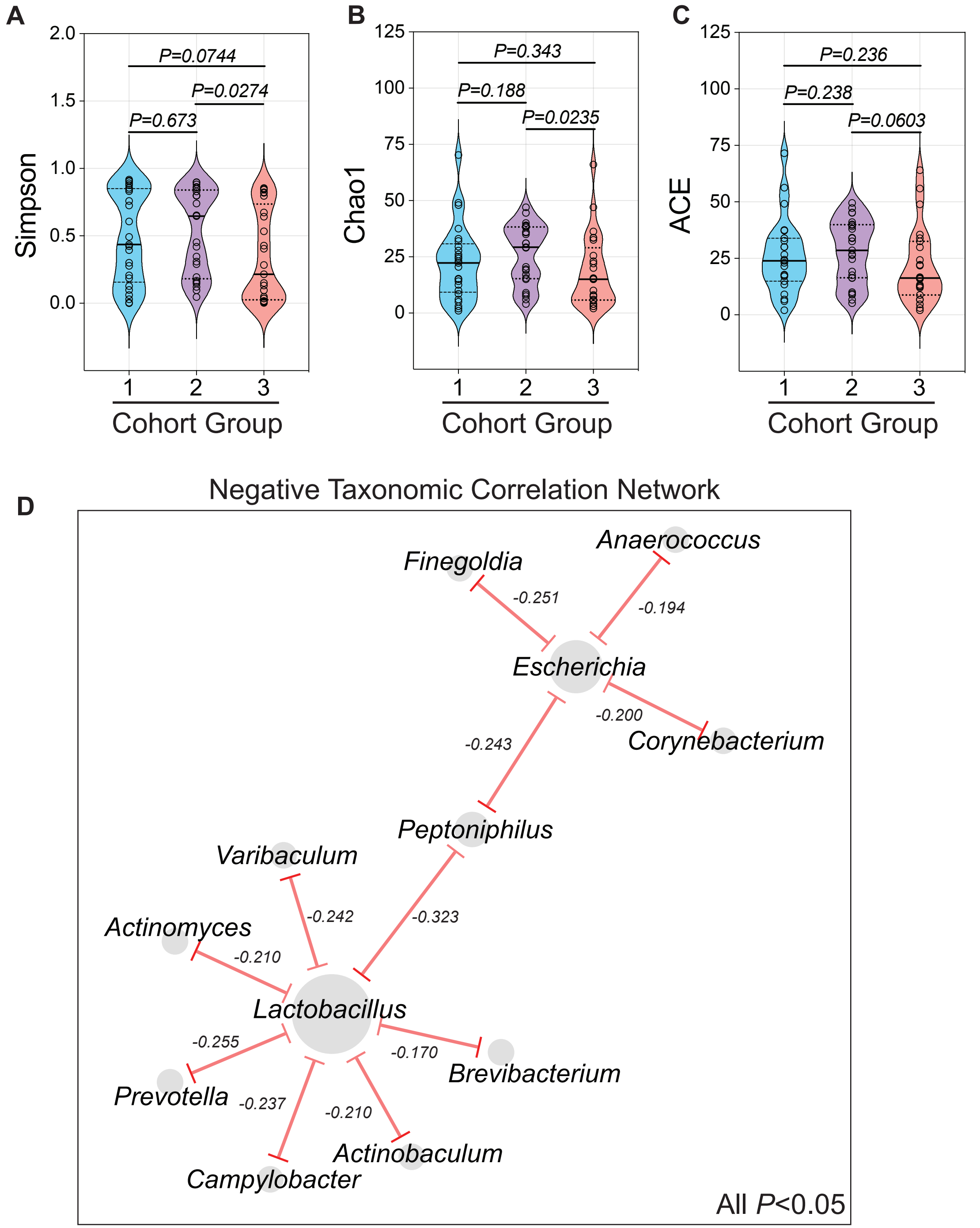

A
2.0
1.5
1.0
0.5
0.0
Simpson
P=0.0744
P=0.0274
P=0.673
1
2
3
Cohort Group
B
125
100
75
50
25
0
Chao1
P=0.343
P=0.188
P=0.0235
1
2
3
Cohort Group
C
125
100
75
50
25
0
ACE
P=0.236
P=0.238
P=0.0603
1
2
3
Cohort Group
D
Negative Taxonomic Correlation Network
Anaerococcus
Finegoldia
-0.251
-0.194
Escherichia
-0.200
Corynebacterium
-0.243
Peptoniphilus
Varibaculum
-0.242
-0.323
Actinomyces
-0.210
Lactobacillus
-0.255
-0.170
Brevibacterium
Prevotella
-0.237
-0.210
Actinobaculum
Campylobacter
All P<0.05

### Figure S6

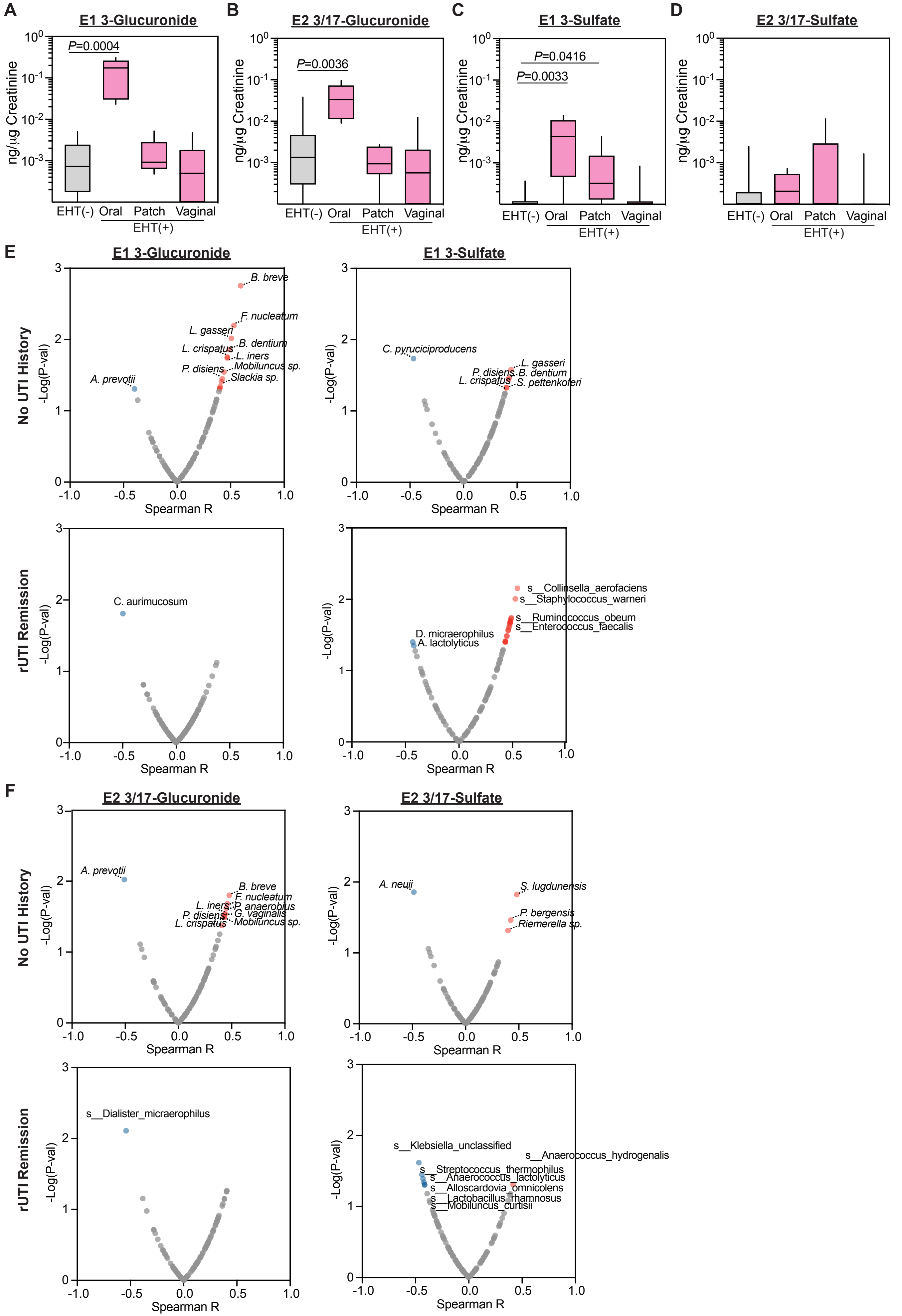
A
E1 3-Glucuronide
P=0.0004
ng/µg Creatinine
EHT(-) Oral Patch Vaginal
EHT(+)
B
E2 3/17-Glucuronide
P=0.0036
C
E1 3-Sulfate
P=0.0416
P=0.0033
D
E2 3/17-Sulfate
E
E1 3-Glucuronide
E1 3-Sulfate
No UTI History
rUTI Remission
-Log(P-val)
Spearman R
B. breve
F. nucleatum
L. gasseri
B. dentium
L. crispatus
L. iners
P. disiens
Mobiluncus sp.
Slackia sp.
A. prevotii
C. pyruciciproducens
S. pettenkoferi
C. aurimucosum
s__Collinsella_aerofaciens
s__Staphylococcus_warneri
s__Ruminococcus_obeum
s__Enterococcus_faecalis
D. micraerophilus
A. lactolyticus
F
E2 3/17-Glucuronide
E2 3/17-Sulfate
P. anaerobius
G. vaginalis
A. neuii
S. lugdunensis
P. bergensis
Riemerella sp.
s__Dialister_micraerophilus
s__Klebsiella_unclassified
s__Anaerococcus_hydrogenalis
s__Streptococcus_thermophilus
s__Anaerococcus_lactolyticus
s__Alloscardovia_omnicolens
s__Lactobacillus_rhamnosus
s__Mobiluncus_curtisii

**Table S1. Cohort clinical features.** Distribution of clinical features among the three groups. Medians and 95% confidence interval are presented for Age, BMI, Urine pH, and Urinary creatinine. *P*-values generated by Kruskal-Wallis test or $\chi^2$ test. Significant differences between groups have been bolded.

**Figure S1. Power analysis and metagenomic dataset characteristics.**
(A) ANOVA power analysis between 3 groups with an a of 0.05 for a range of sample sizes per group. Red lines represent small medium and large effect sized. (B) Two-way T-test power analysis a of 0.05 for a range of sample sizes per group. Red lines represent small, medium, and large effect sizes (Cohen's d). (C) Metagenomic DNA yields within the three cohort groups and commercially available community standards (Zymo). Bars are drawn from the minimum to maximum of the distribution. Boxed represent the interquartile range. Median is depicted by solid line. (D) Average proportion of human genomic content in the WGMS data among the three cohort groups (E) Correlation of theoretical and observed relative abundance of the ZymoBIOMICS Microbial Community standard (Log Distribution). *P*-value generated by permutation.

**Figure S2.**
(A) Genera- and species-level taxonomic profile of taxa detected in sequenced water controls. (B) Advanced urine culturing coverage of each patient among the three cohort groups.

**Figure S3. Taxonomic profiles of detected Archea, Eukaryota, and Vial species.**
(A) Species-level taxonomic profile of Archea and Eukaryota among cohort groups (No UTI History (n=25), rUTI Remission (n=25), rUTI Relapse (n=25)). (B) Species-level taxonomic profile of Viruses among cohort groups (No UTI History (n=25), rUTI Remission (n=25), rUTI Relapse (n=25)).

**Figure S4. Ecological modeling indices among the cohort groups.**
(A) Simpson index, (B) Chao1 index, (C) and ACE index comparison between the three cohort groups (No UTI History (n=25), rUTI Remission (n=25), rUTI Relapse (n=25)). P-value was generated by Kruskal-Wallis test with uncorrected Dunn's multiple correction post hoc. (D) Network analysis of all genera anticorrelation associations with *P*-value less than 0.05. Nodes represent genera edges are defined by Pearson correlation. Node size is proportional to the degree of the node.

**Figure S5. Urinary estrogen conjugate concentrations and taxonomic associations.** Creatinine (Cr)-normalized urinary E1 3-Gluccuronide (A) E2 3/17-Glucuronide (B) E1 3-Sulfate (C) and E2 3/17-Sulfate (D) measured in the urine of EHT(-) and EHT(+) women from the No UTI History and rUTI Remission groups striated by EHT modality (Oral (n=6), Patch (n=6), Vaginal (n=17)). Error bars are drawn from minimum to maximum of the data distribution. Boxes represent the interquartile range. Solid lines denote the median. P-value generated by Kruskal-Wallis test with uncorrected Dunn's multiple correction post hoc. (E) Volcano plots depicting correlation of bacterial species with summed Cr-normalized urinary E1 (E) and E2 (F) conjugates by Spearman correlation in No UTI History (Top panels) and rUTI remission (Bottom panels) groups. P-value generated by permutation. Red dots represent significant ($p<0.05$) positive associations. Blue dots represent significant negative associations.
